# Accurate quantification of single-cell and single-nucleus RNA-seq transcripts using distinguishing flanking k-mers

**DOI:** 10.1101/2022.12.02.518832

**Authors:** Kristján Eldjárn Hjörleifsson, Delaney K. Sullivan, Nikhila P. Swarna, Guillaume Holley, Páll Melsted, Lior Pachter

## Abstract

In single-cell and single-nucleus RNA sequencing, the coexistence of nascent (unprocessed) and mature (processed) mRNA poses challenges in accurate read mapping and the interpretation of count matrices. The traditional transcriptome reference, defining the ‘region of interest’ in bulk RNA-seq, restricts its focus to mature mRNA transcripts. This restriction leads to two problems: reads originating outside of the ‘region of interest’ are prone to mismapping within this region, and additionally, such external reads cannot be matched to specific transcript targets. Expanding the ‘region of interest’ to encompass both nascent and mature mRNA transcript targets provides a more comprehensive framework for RNA-seq analysis. Here, we introduce the concept of distinguishing flanking *k*-mers (DFKs) to improve mapping of sequencing reads. We have developed an algorithm to identify DFKs, which serve as a sophisticated ‘background filter’, enhancing the accuracy of mRNA quantification. This dual strategy of an expanded region of interest coupled with the use of DFKs enhances the precision in quantifying both mature and nascent mRNA molecules, as well as in delineating reads of ambiguous status.

## Introduction

The utility of single-cell RNA-seq measurements for defining cell types has represented a marked improvement over bulk RNA-seq, and has driven rapid development and adoption of single-cell RNA-seq assays (Zeng 2022). One application of single-cell RNA-seq that is not possible with bulk RNA-seq is the study of cell transitions and transcription dynamics, even via snapshot single-cell RNA-seq experiments (Gorin et al. 2022; Gorin, Vastola, and Pachter 2023; La Manno et al. 2018). Such applications of single-cell RNA-seq are based on the quantification of both unprocessed and processed mRNAs (**Supplementary Figure 1**), lending import to the computational problem of accurately and separately quantifying these two modalities (Soneson et al. 2021). The importance of quantifying unprocessed mRNAs in addition to processed mRNAs has also been brought to the fore with single-nucleus RNA-seq (Kuo, Hansen, and Hicks 2023; Grindberg et al. 2013; Ding et al. 2020).

The traditional approach in quantifying RNA-seq has been to rely on a reference transcriptome that defines a ‘region of interest’—typically restricted to mature mRNA transcripts (i.e. no introns) for bulk RNA-seq analyses. This conventional focus has been adequate for the broad objectives of bulk sequencing but is insufficient for the more granular and precise requirements of single-cell and single-nucleus RNA-seq. In these more detailed analyses, reads that emanate from outside the traditional ‘region of interest’ present two problems: a high risk of mismapping within this defined region and the inability to be matched to specific targets within the transcriptome index.

To address the problem of mismapping of external reads (Kaminow, Yunusov, and Dobin 2021), we introduce **distinguishing flanking *k*-mers (DFKs)** (**Figure 1, Algorithm 1, Methods**) to identify reads that are external to the sequences present in the transcriptome index. DFKs are a minimal set of *k*-mers that can be used to distinguish whether a read that is mapped to a set of targets in the transcriptome index has its origin from within the transcriptome index or has an external origin. These *k*-mers thus can act as a filter to prevent reads of external origin from being mismapped to the transcriptome index. In other words, these *k*-mers, if present in a read, will cause the read to be filtered out. We use the term **D-list** to denote the sequences from which DFKs are extracted based on the contents of the transcriptome index. By default, the D-list is set to the genome FASTA file. Therefore, hereinafter, specifying the usage of a D-list refers to supplying the genome FASTA file as the D-list. While using standard mature mRNA transcriptome index with a D-list can be used to improve the quantification of single-cell RNA-seq due to intronic and intergenic reads, still, only mRNA transcripts exist in the index and hence, only reads mapping to mature mRNA regions will be considered. While this is useful for certain applications of single-cell analyses such as cell type identification, having only a single-cell count matrix prevents the usage of biophysical models which jointly consider mature and nascent RNA quantifications (Gorin and Pachter 2023b; Carilli et al. 2023). Thus, extending the index (Melsted et al. 2021; He et al. 2022; Soneson et al. 2021) to allow quantifications of RNA molecules at different stages of their processing is important, as will be discussed next.

**Figure 1.**
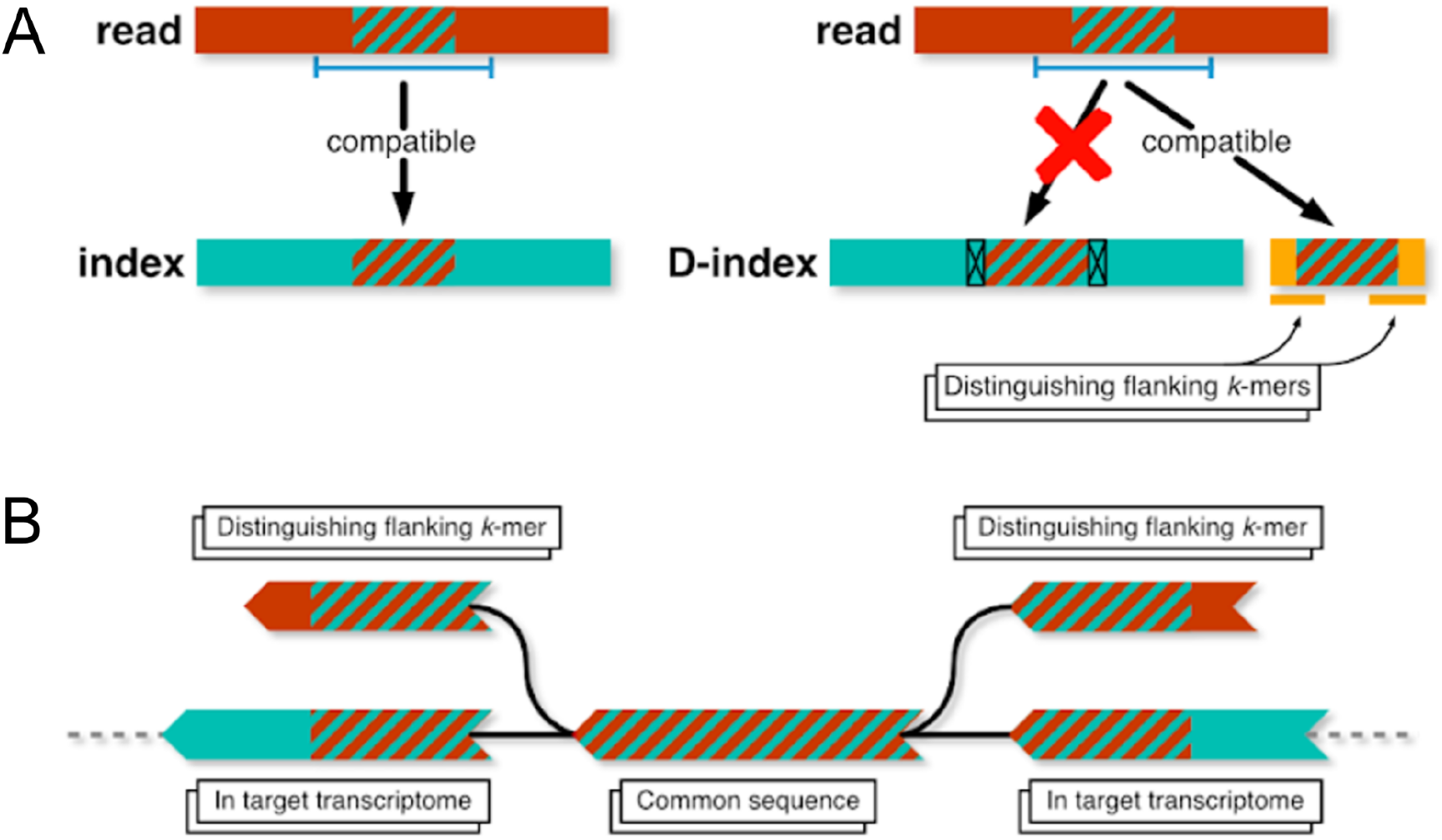
Overview of Distinguishing Flanking *k*-mers. **A)** A non-transcriptomic read containing a subsequence of length greater than *k*, which also occurs in a transcript in the target transcriptome index will get attributed to that transcript. Distinguishing flanking *k*-mers (DFKs), here shown in the modified index (the D-index), can be used to determine whether a read compatible with a reference transcriptome may have originated from elsewhere in the genome. In this diagram, the D-index is one constructed with DFKs. **B)** A de Bruijn graph representation of DFKs.

To quantify nascent RNA transcripts, it is necessary to extend the transcriptome index to include such targets. That is, an index should be created that encompasses the nascent RNAs and the mature RNAs. While seemingly straightforward to construct such an index and to map reads against it, a difficulty arises from classifying individual reads, or individual unique molecular identifiers (UMIs), as being of “mature” or “nascent” status. This difficulty stems from the fact that sequenced reads are typically much shorter than transcripts, and therefore there can be ambiguity in classification of reads as “mature” or “nascent”. Reads that span an exon-exon junction must originate from a completely or partially processed mRNA (which we call “mature”), whereas reads containing sequence unique to an intron must originate from a completely unprocessed or partially processed mRNA (which we call “nascent”). However there are many reads for which it is impossible to know whether they originated from an unprocessed or processed transcript (hence, are ambiguous) (**Figure 2**). Methods that rely on *k*-mer mapping must account for the distinction between *k*-mer ambiguity and read ambiguity, and this distinction has not been carefully accounted for in previous *k*-mer based single-cell RNA-seq pre-processing workflows (Melsted et al. 2021; He et al. 2022).

**Figure 2.**
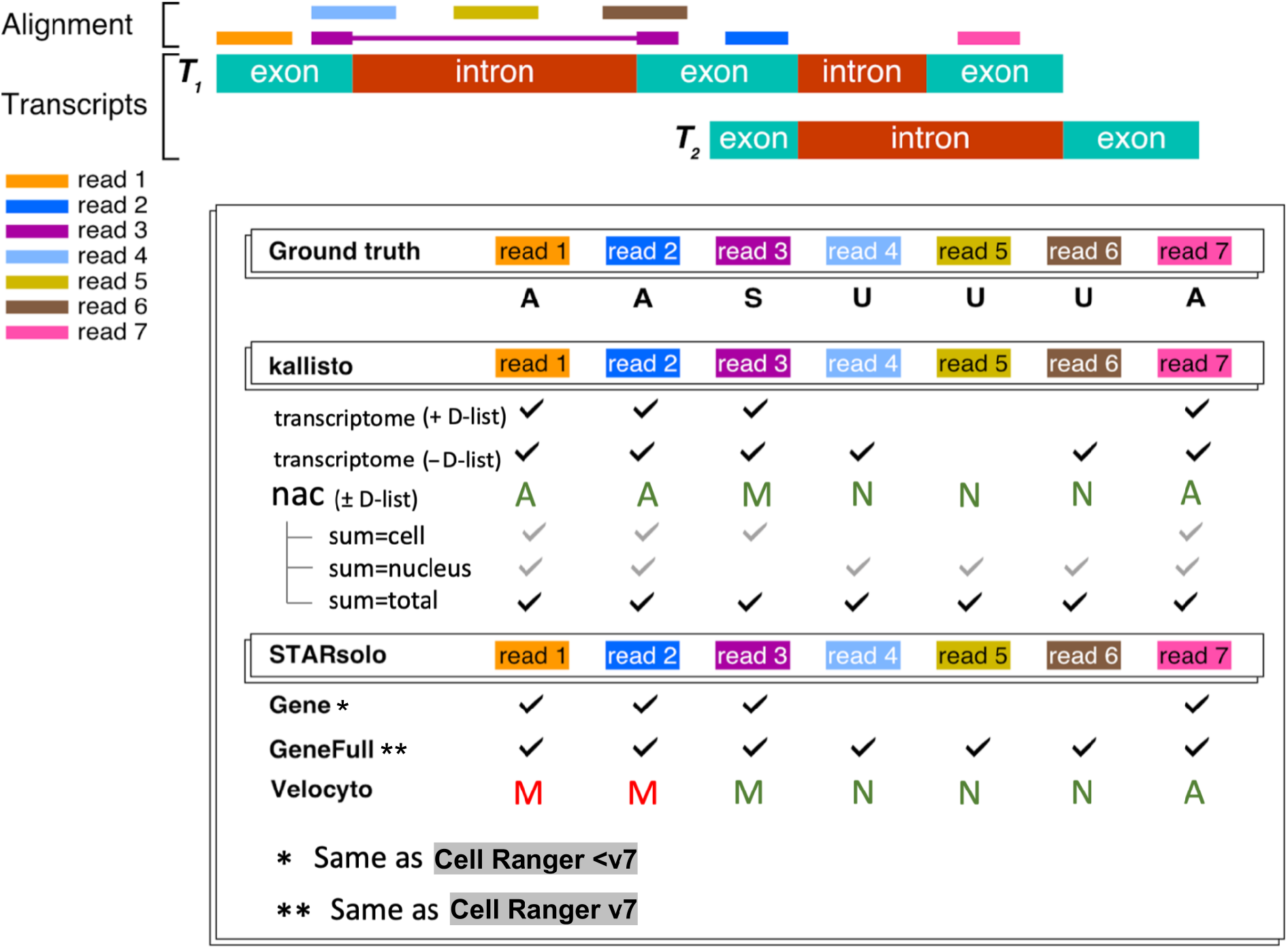
Approach to read assignment and classification into nascent, mature, and ambiguous categories by kallisto, STARsolo, and Cell Ranger. This classification of reads enables accurate classification of RNA species, enabling ambiguous (A) reads to be assigned in various ways based on context (e.g. ambiguous reads are allocated to “mature” (M) in single-cell RNA-seq splicing analysis or added to both “mature” (M) and “nascent” (N) in the case of quantifying “total” RNA content). The kallisto nac index in this example produces the same classification with or without the D-list because no external reads (i.e. those existing outside annotated genomic regions) are present. Results for alevin-fry are not shown because its spliceu index produces classifications identical to kallisto. The checkmarks represent whether a given read will be counted and the letters M, N, and A represent the read classifications (with red letters denoting classifications that differ from the ground truth).

To classify reads as nascent, mature, or ambiguous, we first pseudoalign reads using kallisto (Bray et al. 2016) against a kallisto index containing the mature mRNA (as used originally in pseudoalignment) and nascent mRNA. The nascent mRNA spans the full length of a gene and contains both the gene’s exons and introns as a single contiguous sequence. This comprehensive representation, implemented as the **nac index** in kallisto (Sullivan et al. 2023), allows for accurate classification of reads as mature or nascent or ambiguous, as it properly accounts for the exon-intron boundary and acknowledges that exons are components of both nascent and mature mRNA (**Figure 2**). The developers of alevin-fry have also adopted this approach in the alevin-fry *spliceu* index (He, Soneson, and Patro 2023), an updated version of the original alevin-fry *splici* index, in response to us. However, other tools, such as STARsolo (Kaminow, Yunusov, and Dobin 2021) and the popular Cell Ranger software (Zheng et al. 2017), do not produce such classifications.

While the nac index contains both mature and nascent mRNA, reads of external origin could still arise from intergenic regions of the genome being sequenced. These reads may still be erroneously mapped to this extended transcriptome index. To mitigate the possibility of such instances occurring, one would want to use DFKs by using the nac index with a D-list. This approach is implemented in kallisto by default when building the nac index. Altogether, the nac index, in conjunction with DFKs to mask out reads of external origin, enables the accurate quantification and classification of nascent, mature, and ambiguous mRNA.

## Results

### Distinguishing flanking *k*-mers improve single-cell RNA-seq quantification based on simulations

To assess improvement of using pseudoalignment with distinguishing flanking *k*-mers (DFKs) on single-cell RNA-seq reads, we used the simulation framework developed by the authors of STARsolo (Kaminow, Yunusov, and Dobin 2021). In that simulation framework, errors were introduced into reads at 0.5% mismatch error rate, and reads were simulated from both coding and non-coding genomic sequence to mimic the presence of both unprocessed, partially processed, and completely processed transcripts in single-cell RNA-seq experiments. The top 5000 barcodes, based on UMI count from the simulated data, were used for analysis (**Figure 3A**). When quantifying simulated reads that only span exons with kallisto, the DFKs produced by a D-list do not considerably affect quantification accuracy (**Figure 3B**). However, upon including reads that span introns, the D-list improves the concordance between kallisto quantification count matrix and the simulated truth count matrix in both simulations that only include reads that map uniquely to one gene (**Figure 3C**) and in simulations that additionally include multi-gene reads (**Figure 3D**). Interestingly, although the nac index type includes nascent and mature transcripts, the quantification accuracy still improves slightly with the use of a D-list, likely due to filtering out reads that originate from outside annotated genic loci. The evaluation metrics are shown in **Table 1**. Note that, for the nac index type, UMIs assigned to nascent transcripts were not used in the quantification because the simulation truth matrix does not include nascent transcript counts. For the multi-gene case, bustools was run with the multimapping option enabled when counting UMIs following kallisto quantification. Disabling this option resulted in slightly worse results, due to more false negatives, on the multi-gene simulation (**Supplementary Figure 2**). Enabling the multimapping mode had little effect when run on the exon-only simulations or the non-multi-gene simulations (**Supplementary Figure 3**). While this mode can identify non-uniquely-mapped reads by dividing UMI counts uniformly amongst the genes that the UMI is assigned to, it results in counts that are not whole numbers; thus, the standard for the field has been to discard such UMIs. Finally, since each DFK is only one *k*-mer flanking a unitig, we sought to assess whether considering more *k*-mers flanking a unitig as DFKs (i.e. longer overhangs) would improve the accuracy of kallisto (**Figure 4A**). We found that the benefit of including longer overhangs is negligible (**Figure 4B, Figure 4C, Supplementary Table 1**) therefore, by default, we adhere to having exactly one DFK overhang.

**Table 1.**
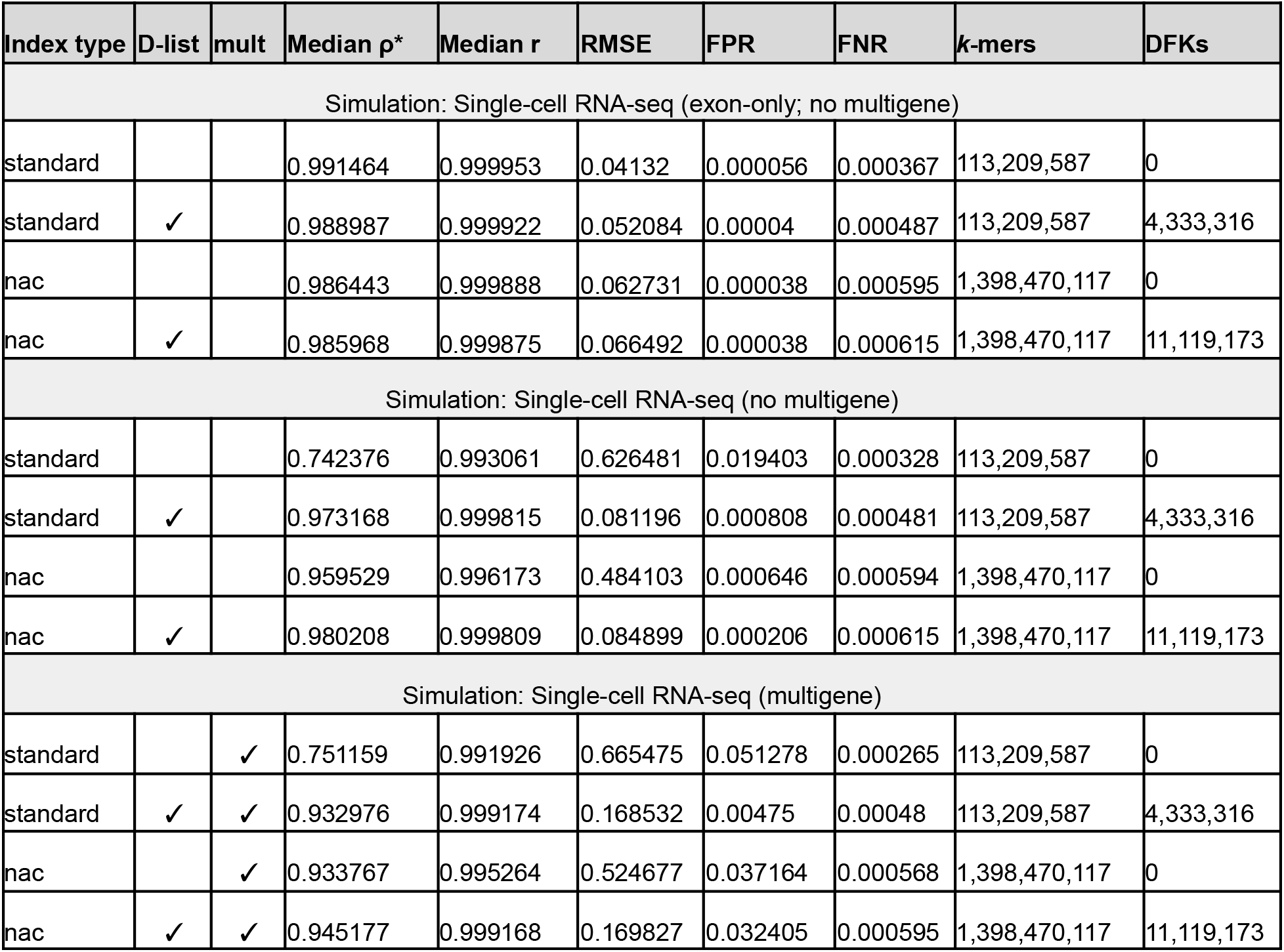
Evaluation metrics of kallisto on simulated data generated using the STARsolo simulation framework. mult: the multimapping quantification mode is enabled. ρ*: Modified spearman correlation. r: Pearson correlation. RMSE: root mean squared error. FPR: false positive rate. FNR: false negative rate. See **Methods** for details.

**Figure 3.**
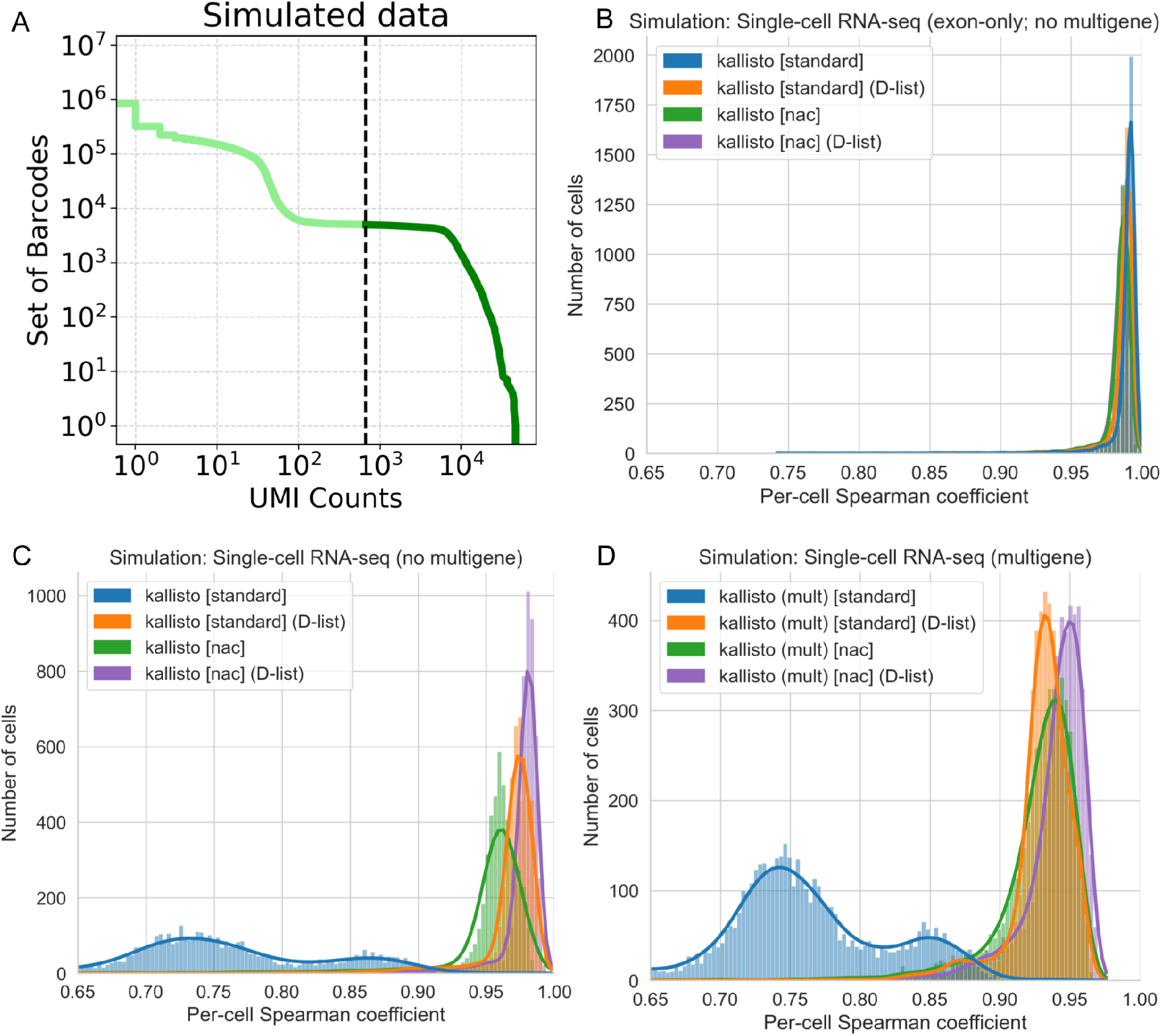
Assessment of the impact of distinguishing flanking *k*-mers on accuracy when tested on simulated data generated using the STARsolo simulation framework.. **A)** Knee plot of the truth count matrix from the STARsolo single-cell RNA-seq simulation. This simulated data represents the “no multigene” simulation. The 5000 cell barcodes with the highest UMI counts were filtered for (corresponding to a UMI threshold of 667). These 5000 cell barcodes were used in downstream analysis of all STARsolo simulated data. **B)** Correlation between kallisto quantifications versus simulated truth for reads only spanning exons. **C)** Correlation between kallisto quantifications versus simulated truth for single-cell RNA-seq reads that map to a single gene. **D)** Correlation between kallisto quantifications versus simulated truth for single-cell RNA-seq reads that include multigene reads. mult: the multimapping quantification mode is enabled. The per-cell spearman correlation, ρ*, between gene counts was determined by excluding genes that contain zero counts in both the kallisto quantification and in the simulation quantification for a given cell barcode (see **Methods**).

**Figure 4.**
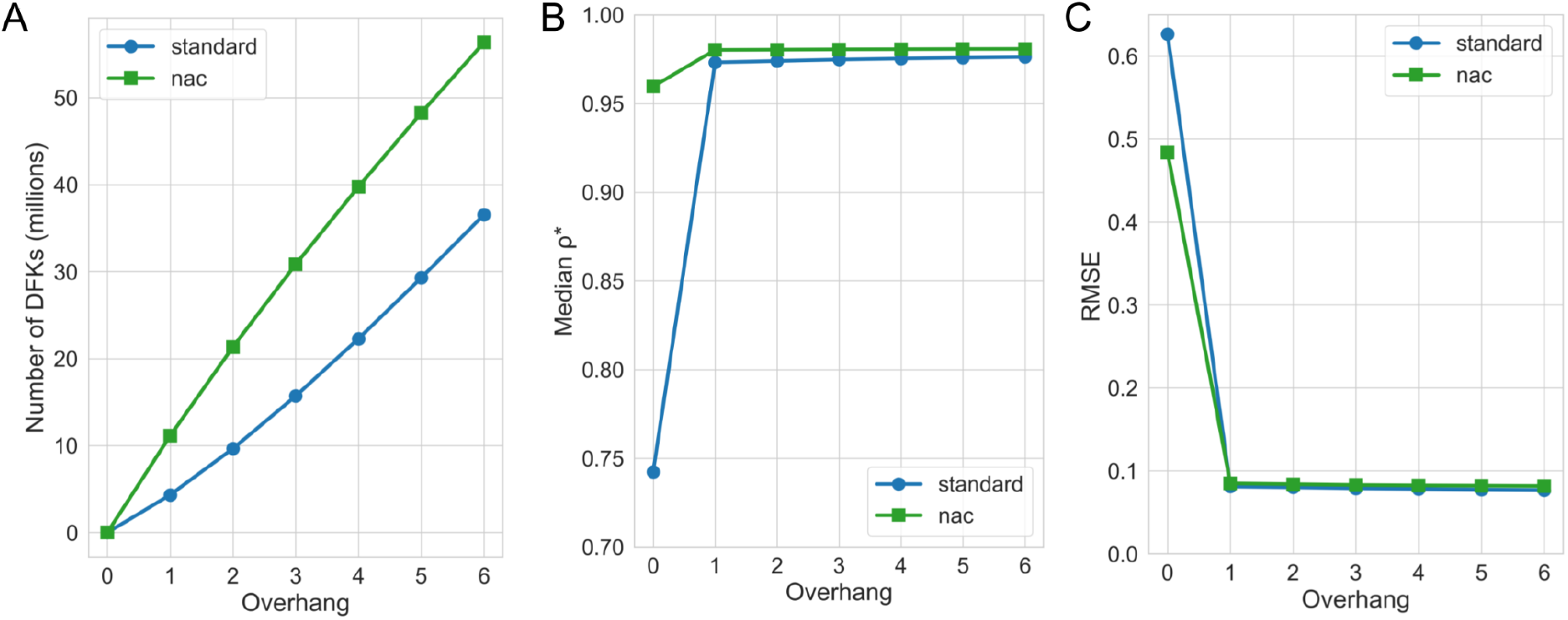
Assessment of the impact of longer overhang distinguishing flanking *k*-mers on accuracy when tested on simulated data generated using the STARsolo simulation framework. **A)** The number of distinguishing flanking *k*-mers (DFKs) at various overhang settings. An overhang of 0 means no DFKs were used. An overhang of 1 is the default setting for the D-list implementation. **B)** Median correlation coefficient ρ* (see **Methods**) between kallisto quantifications at various D-list overhang settings versus simulated truth for the “single-cell RNA-seq (no multigene)” simulation. **C)** RMSE (see **Methods**) between kallisto quantifications at various D-list overhang settings versus simulated truth for the “single-cell RNA-seq (no multigene)” simulation.

### Assessment of other pseudoalignment and alignment based single-cell RNA-seq workflows on simulations

Next, we assessed the performance of other tools using the STARsolo simulation framework. Specifically, we assessed four tools: 1) STARsolo (Kaminow, Yunusov, and Dobin 2021), a single-cell/nucleus RNA-seq tool built into the STAR aligner (Dobin et al. 2013) program, 2) Cell Ranger (Zheng et al. 2017), the pipeline implemented by 10X genomics, 3) cellCounts(Liao et al. 2023), a tool based on the Rsubread aligner (Liao, Smyth, and Shi 2019) and the featureCounts (Liao, Smyth, and Shi 2014) program, and 4) alevin-fry (He et al. 2022; He and Patro 2023), a tool that leverages salmon (Patro et al. 2017) for pseudoalignment. We found that the tools produced quantifications that correlated well with the simulated ground truth for the simulated reads that only span exons (**Figure 5A, Supplementary Table 2**). However, on simulations including intronic reads, both alevin-fry, when executed in a standard pseudoalignment configuration against a spliced transcriptome, and cellCounts performed less well compared to STARsolo and Cell Ranger (**Figure 5B, Supplementary Table 2**). In the case of alevin-fry, using an expanded index that includes introns eliminated this decrease in performance, which is consistent with prior reports (He et al. 2022). Additionally, enabling selective alignment mode (Srivastava et al. 2020) in alevin-fry resulted in further accuracy improvements, similar to the improvements yielded by the D-list, even when used with an expanded transcriptome index.

**Figure 5.**
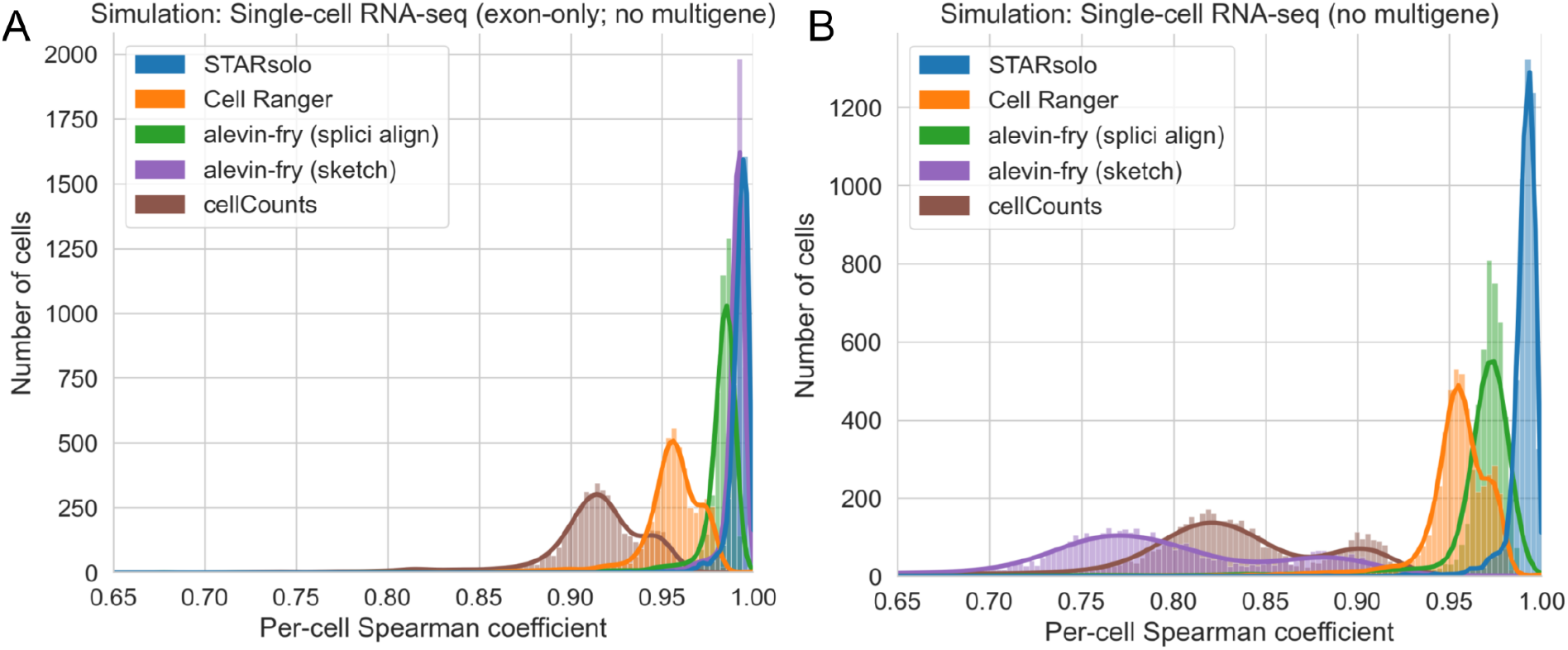
Assessment of different tools on simulated data generated using the STARsolo simulation framework. **A)** Correlation between quantifications produced by the tools versus simulated truth for reads only spanning exons. **B)** Correlation between quantifications produced by the tools versus simulated truth for single-cell RNA-seq reads that map to a single gene. Evaluation against multi-gene reads was not performed because of different methods exposed by different tools to handle such reads. The per-cell spearman correlation, ρ*, between gene counts was determined by excluding genes that contain zero counts in both the kallisto quantification and in the simulation quantification for a given cell barcode (see **Methods**). splici align: Enabling the index used by alevin-fry that contains introns as well as selective alignment mode. sketch: Selective alignment disabled and index is a standard transcriptome index that does not include introns in alevin-fry. For Cell Ranger, version 7 was used with the include-introns option set to false in order to mimic the default behavior of older versions of Cell Ranger. For cellCounts, the featureType option was set to “exon” (which is the default option) rather than “gene” in order to exclude intronic read quantification.

### Distinguishing flanking *k*-mers incur an only minor increase in memory usage and runtime when mapping RNA-seq reads

We assessed the impact of DFKs on memory usage and runtime when processing RNA-seq reads. Across single-cell and single-nucleus RNA-seq datasets from human and mouse tissue, DFKs resulted in only a minor increase in memory usage and runtime. Memory usage increased by less than 2% which is on the order of megabytes while runtime increased by less than 15% (**Figure 6**). On the other hand, mapping RNA-seq reads with the nac index type resulted in a much more substantial increase in memory usage and runtime compared to the standard index type. These results make sense as the nac index type is 10 times larger than (i.e. contains 10 times as many *k*-mers as) the standard index type, whereas the DFKs extracted from a D-list are only a small percentage (i.e. less than 5%) of the total number of *k*-mers. Thus, DFKs can substantially improve RNA-seq mapping accuracy without having a major impact on performance.

**Figure 6.**
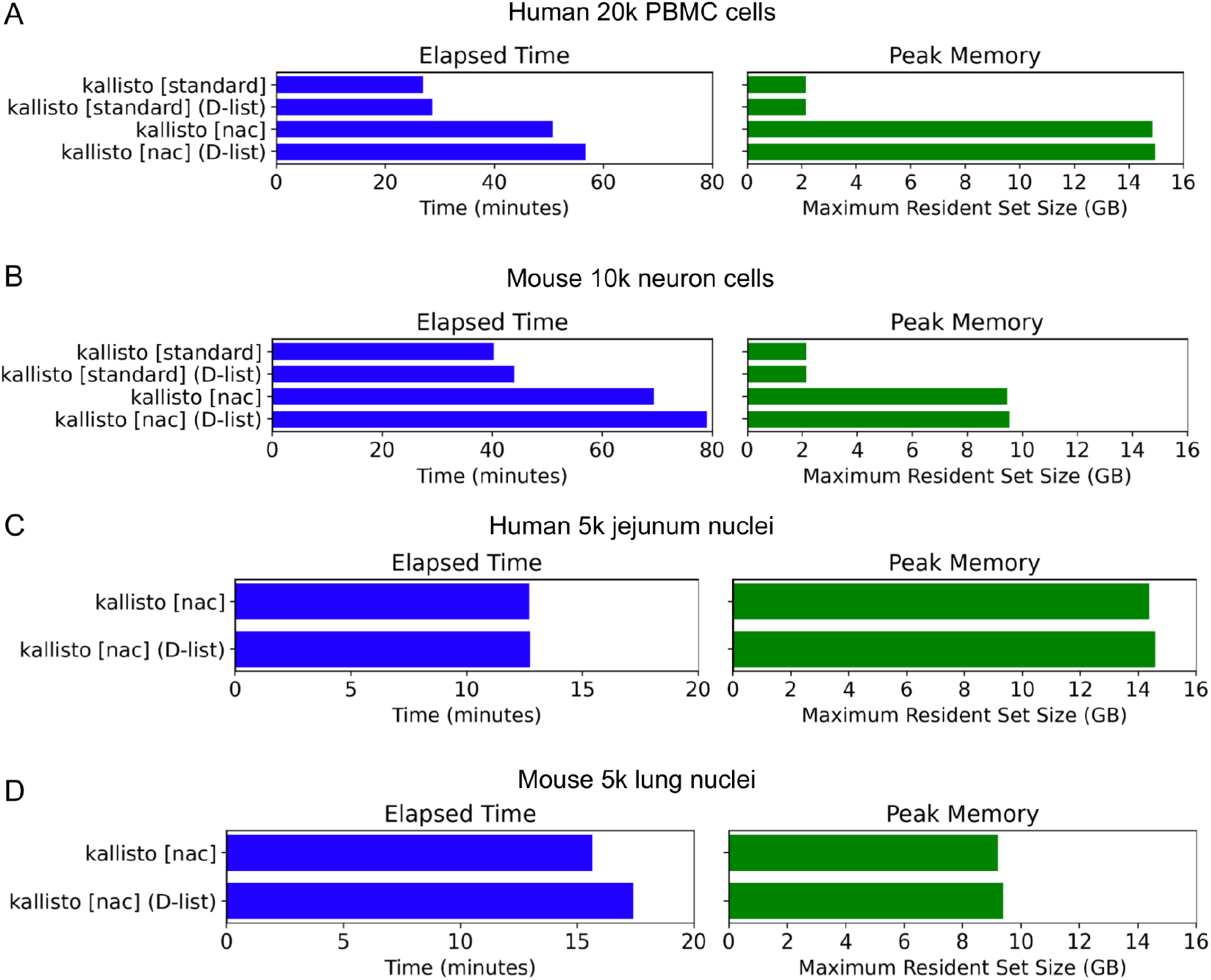
Runtime and memory usage of kallisto with different index types. **A, B)** Runtime and memory usage of the standard index type and the nac index type, created with and without a D-list, on single-cell RNA-seq data. **C, D)** Runtime and memory usage of the nac index type, created with and without a D-list, on single-nucleus RNA-seq data. The standard index type was not employed for single-nucleus RNA-seq data because single-nucleus RNA-seq reads predominantly originate from intron-containing pre-mRNA.

### Distinguishing flanking *k*-mers maintain robustness to sequencing errors during mapping

Since DFKs improve mapping specificity, a natural question that arises is whether the improvement in accuracy scales with higher sequencing error rates. Particularly, how do DFKs compare to alignment-based approaches in maintaining accuracy in the face of more sequencing errors? To address this, we introduced additional sequencing errors, consisting of a combination of mismatches, insertions, and deletions, into the STARsolo simulations (**Supplementary Table 3**). We found that the usage of DFKs always results in an improvement in accuracy, even with a high mismatch rate or a high insertion rate within the simulated sequencing reads (**Figure 7A**). In contrast, while alignment-based methods are very robust to mismatch errors, they fall short with high indel rates (**Figure 7B**). In particular, the same selective alignment settings when executed on the original simulation and simulations where indels are introduced result in a substantial performance decrease on the indel simulations. Altogether, these results suggest that pseudoalignment with the incorporation of DFKs is more robust than alignment-based methods to indels. Such considerations may be important when mapping long-read RNA sequencing reads, which are known to have high indel rates (Zhang, Jain, and Aluru 2020; Delahaye and Nicolas 2021).

**Figure 7.**
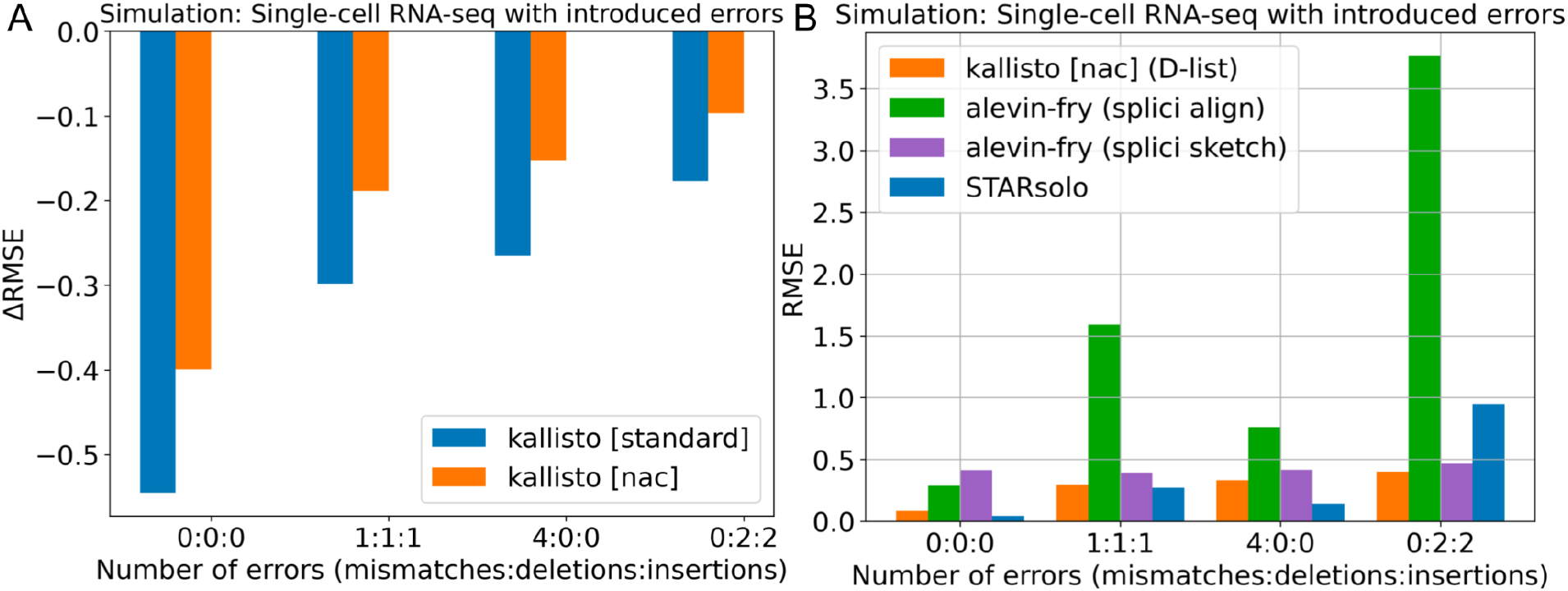
Assessment of different mapping modes on simulated data generated using the STARsolo simulation framework including the introduction of errors into the reads. **A)** Reduction in quantification error, as measured by change in RMSE, by using a D-list to index distinguishing flanking *k*-mers compared to not using a D-list on simulated reads with mismatches, deletions, and insertions. **B)** Quantification error of different tools on simulated reads with mismatches, deletions, and insertions. splici align: Enabling the index used by alevin-fry that contains introns as well as selective alignment mode. splici sketch: Same as “splici align” except selective alignment mode is disabled. The errors were introduced into the “single-cell RNA-seq (no multigene)” simulated reads. RMSE: root mean square error (see **Methods**).

### Nascent, mature, and ambiguous classifications

The quantifications produced by the extended transcriptome index (i.e. the nac index) can classify reads or UMIs as mature (M), nascent (N), or ambiguous (A). We assessed the classifications on both single-cell and single-nucleus data from mouse and human samples. As expected, single-nucleus data tends to have a higher ratio of nascent to mature RNA compared to single-cell data since RNA molecules that have been exported out of the nucleus have undergone splicing and maturation (**Figure 8**) while 10x Genomics Visium spatial transcriptomics data has the lowest proportion of nascent RNA (less than 1%) due to the Visium kit’s exon capture (**Supplementary Figure 4**). Across the different count matrices, we observe that the total counts (N+M+A) are well-correlated with the ambiguous counts, implying that the results of a single-cell or single-nucleus RNA-seq analyses are largely driven by reads mapped solely within exons. Note that regardless of assay type, there tend to be more UMIs classified as nascent than mature, because introns have a much larger coverage over the genome than exon-exon splice junctions. The individual N, M, and A count matrices are poorly correlated with one another, reflecting that different information is present in each of those three matrices. As biophysical models of the RNA life cycle make use of nascent transcript counts and mature transcript counts, how to allocate those ambiguous counts to either nascent or mature remains a topic for future research. For now, one might reasonably assume that the ambiguous counts in single-cell RNA-seq experiments originate from mature transcripts since, in such assays, it is expected that there will be more mature transcripts than nascent transcripts therefore a purely-exonic UMI is most likely to be mature. However, in the nucleus context, the likelihood of a purely-exonic UMI being mature is lower since there will be fewer mature transcripts, as evidenced by the much larger nascent-to-mature ratio in UMI classification. Developing methods to more accurately allocate ambiguous reads is an interesting topic to pursue but is beyond the scope of this present study.

**Figure 8.**
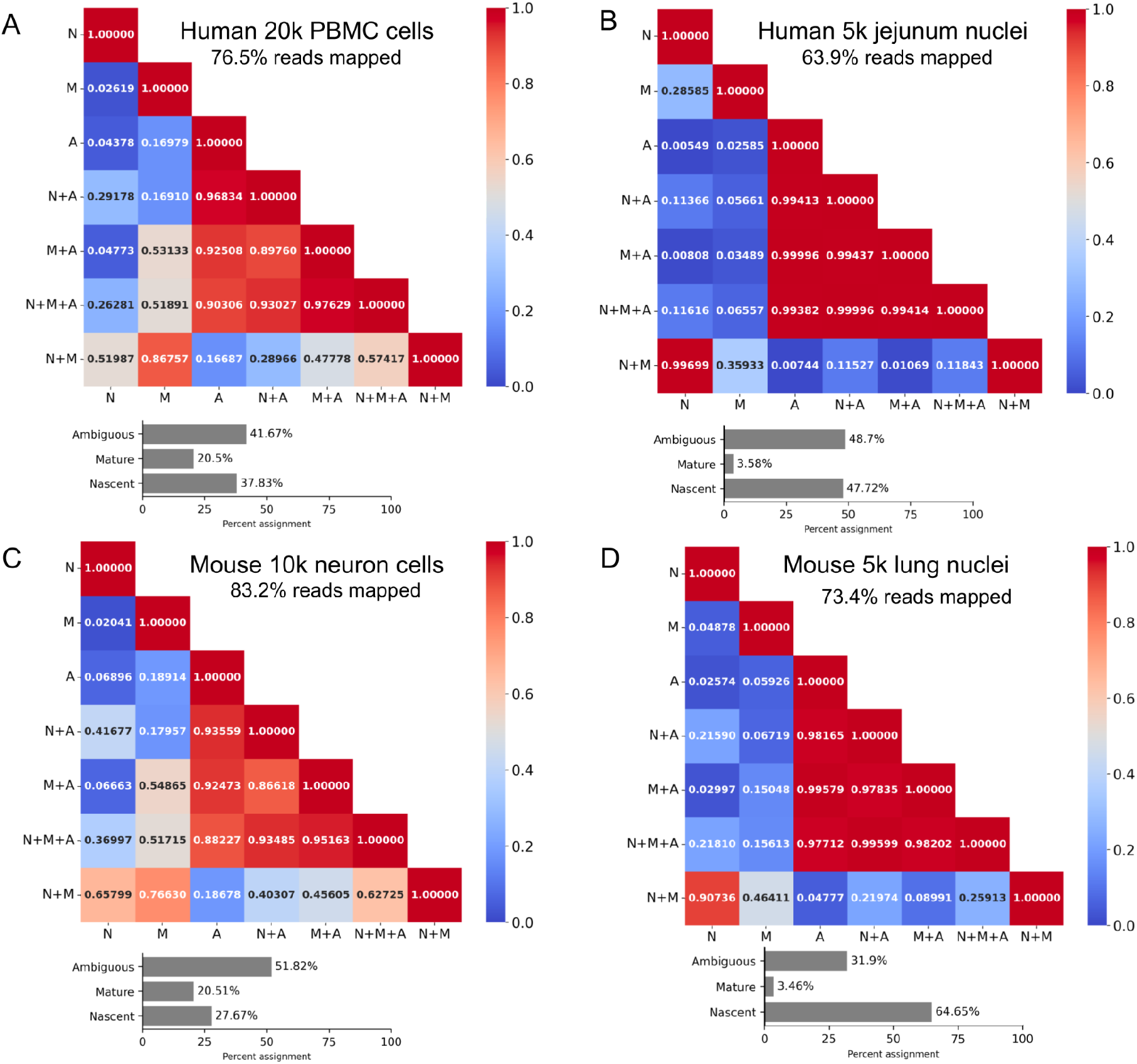
Different count matrices and their combinations produced from single-cell and single-nucleus experiments. **A, B)** Exploration of single-cell and single-nuclei count matrices from human samples. **C, D)** Exploration of single-cell and single-nuclei count matrices from mouse samples. The heatmap shows the Pearson correlation coefficient between pseudo-bulked count matrices. Pseudo-bulking was performed on cell barcodes with at least 500 UMIs detected in the N+M+A (total) matrix. The bar plots show the percentage of UMIs assigned to the ambiguous, nascent, and mature classifications. N: Nascent, M: Mature, A: Ambiguous.

## Discussion

This study introduces a combined approach of using distinguishing flanking k-mers (DFKs) with an extended transcriptome index (nac index) in single-cell and single-nucleus RNA-seq analysis (**Figure 9**). This method aims to address specific challenges in RNA sequencing, particularly in the quantification of nascent and mature mRNA transcripts, and in reducing mismapping errors caused by reads originating outside of the targeted transcriptomic regions.

**Figure 9.**
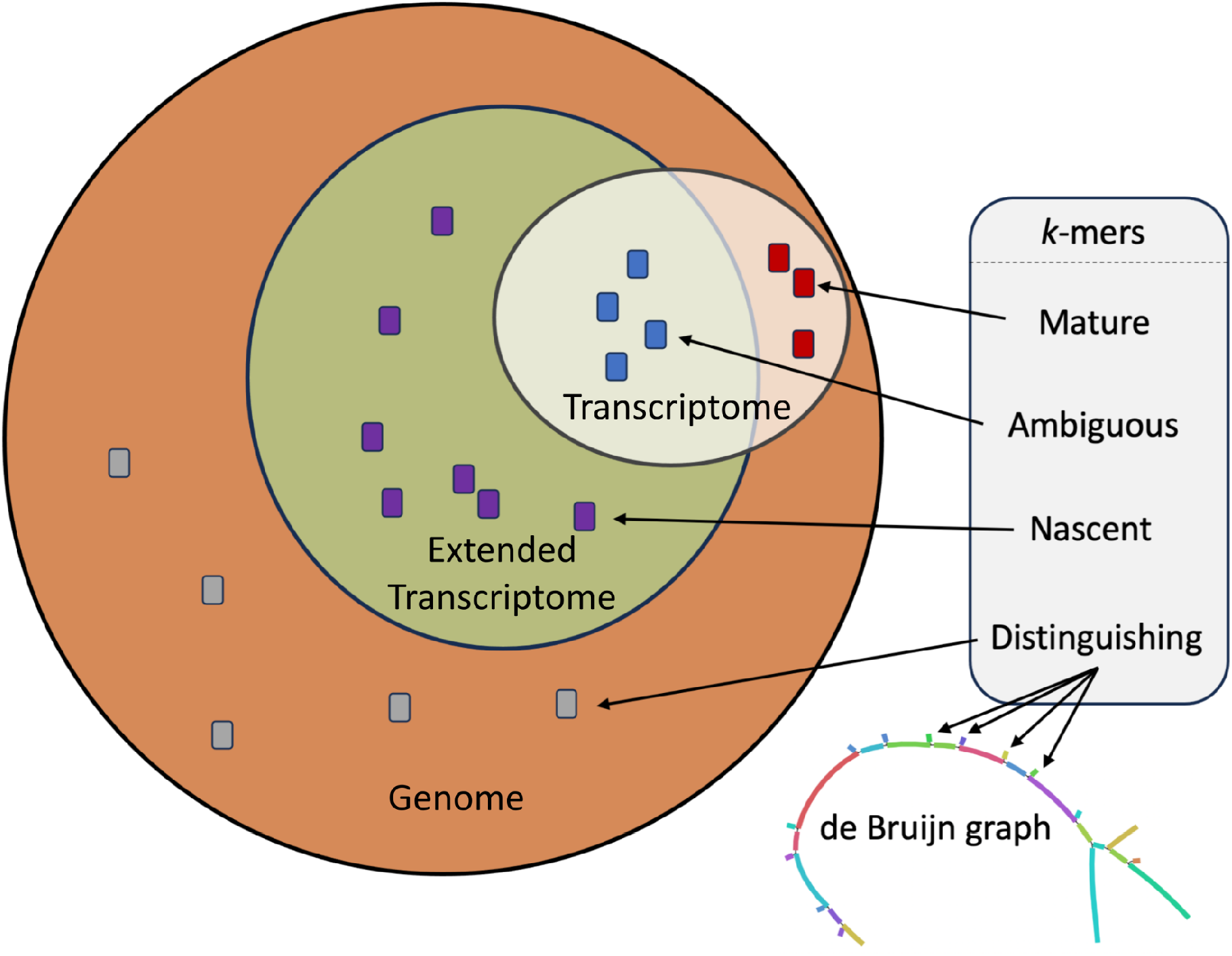
Summary of enhancements to read mapping and classification. This figure shows *k*-mers originating from the standard transcriptome index, the extended transcriptome index containing nascent RNA transcripts, and the entire genome. The integration of distinguishing flanking k-mers into a de Bruijn graph is shown.

While this method can provide accurate quantification of mature RNA transcripts, nascent RNA transcripts, and ambiguous RNA transcripts (i.e. transcripts that cannot be unambiguously resolved as nascent or mature), how to jointly utilize these three types of RNA transcripts remains an avenue for future research. One approach to “integrating” the nascent and mature modalities is via biophysical modeling of transcription (Carilli et al. 2023; Gorin and Pachter 2023b; Gorin, Vastola, and Pachter 2023), however questions remain, such as how to best utilize reads that are ambiguous between the modalities. Importantly, there is not one single “count matrix”; rather, there are multiple count matrices that each lend value in single-cell and single-nucleus RNA-seq analyses. The number of count matrices becomes even larger when considering technologies such as SPLiT-seq (Rosenberg et al. 2018), for which two different priming strategies (oligo-dT and random hexamer) exist for a single cell, or Smart-seq3 (Hagemann-Jensen et al. 2020), for which two cDNA fragment types (UMI and internal) exist, thus resulting in an additional set of count matrices (**Supplementary Figure 5**). The ability to differentiate and quantify nascent, mature, and ambiguous transcripts offers a more nuanced view of gene expression, potentially enriching our understanding of RNA processing and transcriptome dynamics.

There are several limitations to the quantification framework we have proposed. In a cell, the set of unprocessed mRNAs at any given time is likely to include partially processed molecules (Pai et al. 2018; Pandya-Jones and Black 2009), and in principle the complete splicing cascade must be understood and known in order to accurately quantify single-nucleus or single-cell RNA-seq data (Gorin and Pachter 2022). Furthermore, the presence of ambiguous reads both for single-cell and single-nucleus RNA-seq is unsatisfactory. Ideally reads should be longer so that they can be uniquely classified, or they should be fractionally classified probabilistically. The latter approach is non-trivial due to variation in effective transcript lengths that will depend on library preparation and must be accounted for (Gorin and Pachter 2023a; Pachter 2011), but this is an interesting direction of study.

While most mismapping errors that affect single-cell RNA-seq quantification are eliminated by extending the transcriptome index, DFKs provide further improvement to quantification accuracy. Specifically, DFKs can eliminate erroneous mapping of reads that originate from transcripts that appear outside even the extended transcriptome index. More importantly, DFKs provide high scalability. DFKs can scale to higher sequencing error rates as the accuracy gains of DFKs are not reversed when different sequencing error profiles are introduced. Moreover, DFKs can scale to size. When only a small specific set of targets is of interest but there are many known possible target sequences, those possible target sequences can simply be incorporated into the D-list. The resultant DFKs will optimize mapping specificity making it unnecessary to index all the possible target sequences. Irrespective of whether the target sequences occupy a small proportion or a large proportion of the “background”, the DFKs will improve mapping specificity without any major impact on performance. Thus, the DFKs act as a space-efficient general “background filter”.

In summary, this study introduces a method for improving the accuracy of generating count matrices. It is anticipated that these improved quantifications and the multimodal nature of these quantifications will prove useful for multiple downstream applications, including both total gene expression quantification and the integration of multiple count matrices via biophysically informed models.

## Methods

### The D-list

A D-list (distinguishing list) enables accurate quantification of RNA-seq reads in experiments where reads that are not an expression of the target transcriptome may still contain sequences which do occur in the target transcriptome. Without the D-list, these reads may be erroneously quantified as transcripts in the target transcriptome, based on alignment of the common sequences. Thus, the D-list may contain any sequences that are not desired in the abundance matrix yielded by the quantification. Such sequences may include genomes of other organisms (Luebbert et al. 2023) to avoid mismapping due to sample contamination, they may consist of the genome from which the target transcriptome was made, or they may contain common transposable elements, such as Alu regions, which might confound analyses. The D-list is incorporated into the index by finding all sequences, *k* base-pairs or longer, that occur in both the D-list and the target transcriptome. The first *k*-mer upstream and the first *k*-mer downstream of each such common sequence in the D-list are added to the index colored *de Bruijn* graph. We refer to these new vertices in the graph as *distinguishing flanking kmers* (DFKs) (**Algorithm 1**). The DFK vertices are left uncolored in the index, such that during quantification, reads that contain them will be masked out, and go unaligned.

As an illustration, consider a read containing both *k*-mers found only in intergenic RNA and k-mers found both in mRNA and the intergenic RNA. If that read is mapped to an index built from mRNA transcripts, the mRNA *k*-mers will be found in the index, whereas the disambiguating genome *k*-mers will not. The whole read will be erroneously mapped based on the ambiguous *k*-mers (i.e. the k-mers found both in mRNA and the intergenic RNA) that are present in the index. By finding all ambiguous *k*-mers in the mRNA index, and adding any distinguishing flanking genome *k*-mers to the index, the read will be masked from mapping to an mRNA transcript.

Recent papers have discussed various ways of reducing the number of false positives in RNA quantification through either including the entire genome or a subset of the genome in the index as a “decoy” or through alignment scoring (Srivastava et al. 2020). Our method is distinct in that it incorporates only the minimum amount of data, required to disambiguate common sequences, into the index while still adhering strictly to the principles of k-mer based pseudoalignment. Therefore, the memory usage and runtime of using pseudoalignment using a D-list are on par with the memory usage and runtime without the use of a D-list. Our method can scale favorably to larger genome size (or, more generally, larger D-lists) while the target sequences to map against remain small.

The D-list is implemented in kallisto version 0.50.1 (Bray et al. 2016; Sullivan et al. 2023). The kallisto index command contains a --d-list option that takes in, as an argument, the path to a FASTA file containing the D-list sequences for building an index with the D-list. The kallisto index command also contains a --d-list-overhang option for specifying longer overhangs (i.e. extending the flanking sequences that make up the DFKs). The kallisto bus command (Melsted, Ntranos, and Pachter 2019) contains a --dfk-onlist option that, when enabled, adds a D-list target to the equivalence class for a given pseudoalignment if a DFK is encountered rather than discards the read; this option is useful for distinguishing reads that don’t pseudoalign versus reads that are discarded due to a DFK. Finally, in kb-python (version 0.28.0), the kb ref command automatically uses the genome FASTA as the D-list when building the kallisto index – a behavior that can be overwritten by explicitly specifying --d-list in kb ref. As a minor nuance, the default genome FASTA D-list does not contain splice junctions (SJs); however, the number of additional DFKs that would be indexed with the inclusion of SJ-spanning sequences is miniscule since SJ-spanning contigs are only *k* − 1 *k*-mers in length. Therefore, including the spliced transcriptome in the D-list would be unlikely to make any difference in read mapping.

### Generating distinguishing flanking k-mers

A sequence *s* is a string of symbols drawn from an alphabet Σ = {A, T, C, G}. The length of *s* is denoted by |*s*|. A substring of *s* is a string that occurs in *s*: it has (zero-indexed) start position *i* and end position *j* and is denoted by *s*[*i* : *j*], therefore |*s*[*i* : *j*]| is equal to *j* − *i*. In the case that |*s*[*i* : *j*]| equals *k*-mer size *k, s*[*i* : *j*] is *k*-mer. A compact de Bruijn graph (cdBG) is a de Bruijn graph where all maximal non-branching paths of vertices from a de Bruijn graph, wherein each vertex is a *k*-mer, are merged into single vertices (Holley and Melsted 2020). Each vertex in a compact de Bruijn graph is a sequence called a unitig. We define a compact de Bruijn graph *U* as a set where each element *u* ∈ *U* is a unitig. The function Map(*s, u*) takes in a *k*-mer *s* and a unitig *u*, and returns the position of *s* along *u* if *s* exists in *u*, or NULL otherwise. **Algorithm 1** applies these definitions towards identifying distinguishing flanking k-mers (*DFKs*) from a D-list *D*, given a cdBG *U* of *k*-mer size *k* built over target sequences (e.g. a transcriptome). For expository purposes, the algorithm is described such that *U* is a non-bidirected cdBG (i.e. the *k*-mers and their reverse complements are *not* represented identically). However, in practice each *k*-mer and its reverse complement are represented as a single canonical *k*-mer (the lexicographic minimum of the *k*-mer and its reverse complement). Additionally, for simplicity, we define DFKs and describe the algorithm only for single overhangs.

#### Algorithm 1 Generate distinguishing flanking k-mers from a D-list

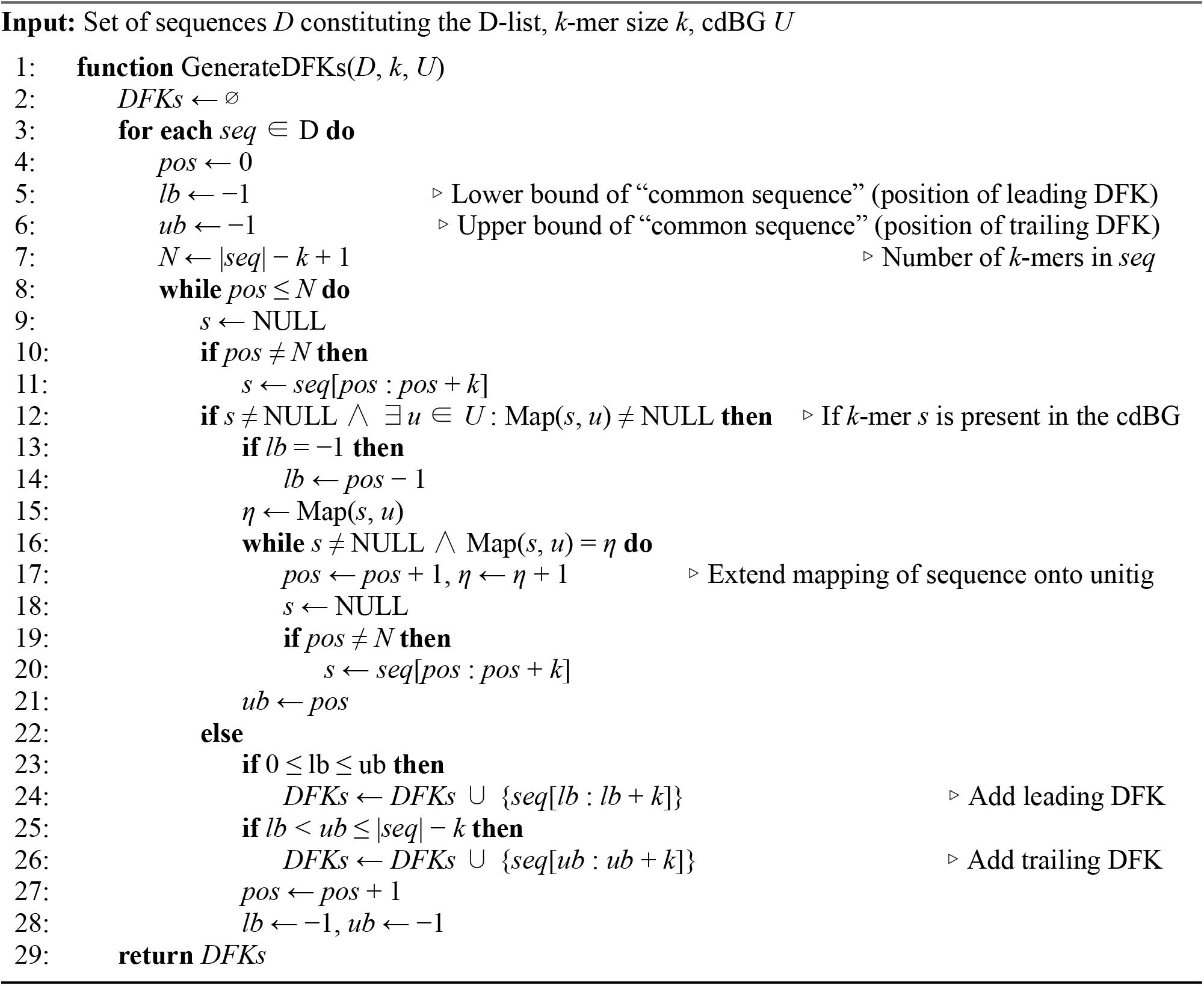

#### Lemma 1.

The worst case space complexity of DFKs is O(min(N_k_, M_k_)) where N_k_ is the number of unique *k*-mers in the de Bruijn graph (dBG) and M_k_ is the number of unique *k*-mers in the D-list.

<u>Proof.</u>

Considering the alphabet Σ = {A,T,C,G}, ∀*s* ∈ dBG, the maximum number of flanking *k*-mers on each side of *s* is |Σ|, permitting a flanking *k*-mer for each character in the alphabet. On each side of *s*, the maximum number of DFKs, which are the flanking *k*-mers in the D-list but not in the dBG, is |Σ| − 1 corresponding to the presence of one flanking *k*-mer that exists in the dBG and the remaining |Σ| − 1 *k*-mers being DFKs. Since *s* has two sides (leading and trailing), the maximum number of DFKs becomes 2(|Σ| − 1) = 6. In the worst-case scenario, ∀*s* ∈ dBG, *s* contains the maximum number of DFKs. Thus, |DFKs| ≤ 6N_k_ where |DFKs| is the cardinality of the set of DFKs. The actual number of DFKs identified from the D-list is bounded by the number of unique *k*-mers in the D-list, denoted as M_k_, i.e., |DFKs| ≤ M_k_. Since |DFKs| ≤ min(6N_k_, M_k_), the space complexity for storing DFKs is O(min(N_k_, M_k_)). ▮

### Improvements to the kallisto index

The *de Bruijn* graph implementation in kallisto was replaced with Bifrost (Holley and Melsted 2020), which employs a minimizer (Roberts et al. 2004) lookup table in lieu of a *k*-mer lookup table in order to achieve a lower memory footprint. Furthermore, since the set of minimizers in the graph is known at the time of quantification, we replaced the minimizer hash function with BBHash (Limasset et al. 2017), which implements a minimal perfect hash function. This enables kallisto to shrink the minimizer hash table to capacity, saving memory. Additionally, equivalence classes (ECs) were restructured. Sets of transcripts are represented as Roaring bitmaps (Chambi et al. 2016) and a Robin Hood hashmap (Leitner-Ankerl 2022) is used for the inverted hash table mapping transcript sets to equivalence class. ECs are allocated dynamically during quantification. Thus, an EC is only created once it has been found to be used by a read, whereas the previous paradigm preemptively created the ECs used by all the vertices in the graph, during indexing, regardless of whether or not they were ever used by a read. Finally, the index includes an external Robin Hood hashmap for storing DFKs since, as the number of DFKs is relatively small, the external hashmap occupies less memory, with only a small reduction in speed, compared to integrating the DFKs into the main de Bruijn graph (**Supplementary Figure 6**). These changes have resulted in an approximately 2x reduction in runtime and 4x reduction in memory consumption in kallisto v0.50.1 compared to kallisto version 0.48.0 when using the nac index type to map single-cell RNA-seq reads (**Supplementary Figure 6**).

### The simulation framework

We obtained the simulation framework developed by the authors of STARsolo (Kaminow, Yunusov, and Dobin 2021) from https://github.com/dobinlab/STARsoloManuscript/ and ran the simulation as-is to generate a ground truth matrix. For the kallisto nac index type, the “mature” and “ambiguous” count matrices were summed up by using --sum=cell in kb count and the resultant matrix was used for testing. For all tools, a predefined “on list” of barcodes (referred to in other tools as a whitelist or an unfiltered permit list) was supplied. The three simulated sequencing datasets used are:

- No multigene: 338,984,644 reads
- With multigene: 350,024,222 reads
- Exon-only, no multigene: 189,451,399 reads

For the simulations where errors were introduced into sequences, the reads were processed to first introduce mismatches, then introduce deletions, then introduce insertions.

### Runtime and memory usage assessment

The command `/usr/bin/time –v` which executes the GNU time program was used to obtain the elapsed (wall clock) time and the maximum resident set size for runtime and peak memory usage, respectively. All performance assessments were conducted on a server with x86-64 architecture, 88 CPUs (Intel Xeon Gold 6152 CPU @ 2.10GHz), and 768 GB of memory.

### Evaluation metrics for single-cell RNA-seq simulations

To evaluate the performance of a program’s output gene count matrix *G*_*p*_ ∈ ℝ^*n×m*^ against a simulation’s ground truth gene count matrix *G*_*s*_ ∈ ℝ ^*n×m*^, where *n* is the number of cells, *m* is the number of genes, *ŷ*_*ij*_ is the count of gene *j* in cell *i* in *G*_*p*_, and *y*_*ij*_ is the count of gene *j* in cell *i* in *G*_*s*_, the following metrics are used:

Root mean squared error:

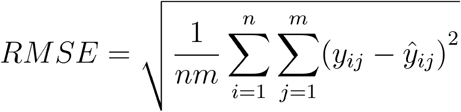

False positive rate:

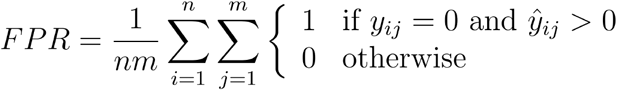

False negative rate:

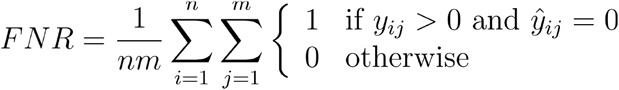

Correlation coefficient:

We use two per-cell correlation coefficients to assess the correlation between the “ground truth” simulated gene counts cell and the program’s output gene counts for a given cell. The first, r, is the Pearson correlation computed across all genes within a given cell. The second, ρ*, is a modified variant of the Spearman correlation in that the Spearman correlation is computed only using the genes that have a non-zero count in both the simulation and the program output within a given cell. This variant is the assessment used by the developers of the STARsolo simulations (Kaminow, Yunusov, and Dobin 2021). The restriction to nonzero cells is necessary when using the Spearman correlation, as the zeroes cannot be ranked with respect to each other. However, we note that use of the Spearman (and therefore ignoring the zeroes) provides an assessment that is highly sensitive to low counts, especially the difference in a program reporting a one or a zero for a gene in a cell.

Pearson correlation for cell *i*, using all genes:

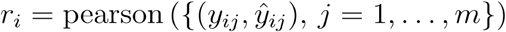

Spearman correlation for cell *i*, using only genes with a non-zero count in both the simulation and the program output for that cell:

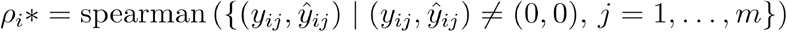

### Datasets used

The human reference genome (GRCh38) used throughout is the same one used in the STARsolo simulations (Kaminow, Yunusov, and Dobin 2021). The GRCh38 FASTA and GTF files used are available from the code repository associated with this paper. The mouse reference genome (GRCm39) used is the primary assembly FASTA file from Ensembl with the corresponding GTF annotation version 110, which was filtered to only include the gene_biotype values of protein_coding, lncRNA, lincRNA, and antisense. The following sequencing datasets, publicly available from 10x genomics (all version 3 chemistry), were used:

- Human 5k PBMC (single-cell): 383,941,607 reads
  ∘ Sample name: 5k_pbmc_v3
- Human 20k PBMC (single-cell): 818,107,363 reads
  ∘ Sample name: 20k_PBMC_3p_HT_nextgem_Chromium_X
- Human 5k jejunum (single-nucleus): 121,378,620 reads
  ∘ Sample name: 5k_human_jejunum_CNIK_3pv3
- Mouse 10k neuron (single-cell): 1,589,915,447 reads
  ∘ Sample name: SC3_v3_NextGem_SI_Neuron_10K
- Mouse 5k lung (single-nucleus): 232,479,932 reads
  ∘ Sample name: 5k_mouse_lung_CNIK_3pv3
- Mouse embryo Visium CytAssist 11mm FFPE: 832,193,962 reads
  ∘ Sample name: CytAssist_11mm_FFPE_Mouse_Embryo

## Code availability

kallisto is available under the BSD-2-Clause license and is available at https://github.com/pachterlab/kallisto. Code for the analyses performed for this paper is available at https://github.com/pachterlab/HSSHMP_2024.

## Software versions

Unless stated otherwise, the software versions used are as follows: kallisto 0.50.1, bustools 0.43.2, kb-python 0.28.0, salmon 1.10.0, alevin-fry 0.8.2, simpleaf 0.15.1, Rsubread 2.12.3, Cell Ranger 7.0.1, STAR 2.7.9a. Additionally, Bandage version 0.8.1 (Wick et al. 2015) was used for rendering de Bruijn graphs into ribbon-like representations.

## Acknowledgments

D.K.S. was funded by the UCLA-Caltech Medical Scientist Training Program (NIH NIGMS training grant T32 GM008042). L.P. was supported in part by the National Institutes of Health (NIH) grants U19MH114830 and 5UM1HG012077-02. Additionally, this work was supported in part by the Icelandic Research Fund Project grant number 218111-051. The development of kallisto and bustools was funded in part by a grant awarded during round 2 of the Essential Open Source Software for Science by the Chan Zuckerberg Initiative for “Open Source Software for Bulk and Single-cell RNA-seq”.

A response to the initial version of the preprint was written (He, Soneson, and Patro 2023), to which we reply in **Supplementary Text 1**.

## Competing interests

The authors declare no competing financial interests.

## Contributions

K.E.H., D.K.S., P.M., and L.P. conceived this study. P.M. and L.P. supervised this study. K.E.H. and D.K.S. implemented the methods described in this study in the kallisto software (version 0.50.1). K.E.H. and D.K.S. integrated the Bifrost de Bruijn graph into the kallisto software with help from G.H. K.E.H., D.K.S., and N.P.S. benchmarked kallisto’s D-list feature. K.E.H. and D.K.S. performed analyses and produced figures. K.E.H., D.K.S., and L.P. contributed to writing the paper. K.E.H. and D.K.S. contributed equally to this work.

## Notes

### Competing Interest Statement

The authors have declared no competing interest.

### Summary of Updates

This version fixes a formatting issue that arose in pdf conversion in v2, and a typo.

