## Supplementary Information for "Accurate quantification of single-cell and single-nucleus RNA-seq transcripts using distinguishing flanking k-mers"

**Supplementary Figure 1**: The maturation process of RNA transcripts.

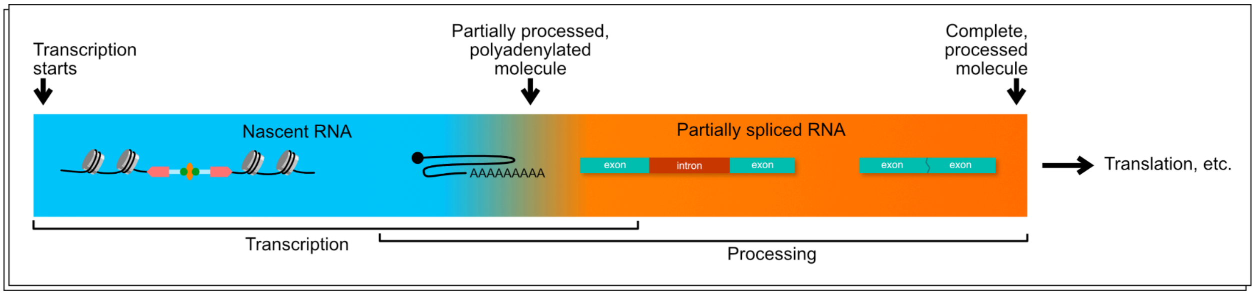

**Supplementary Figure 2**: Assessment of kallisto quantification results, without multimapping mode enabled, on simulated single-cell RNA-seq reads that include multi-gene reads.

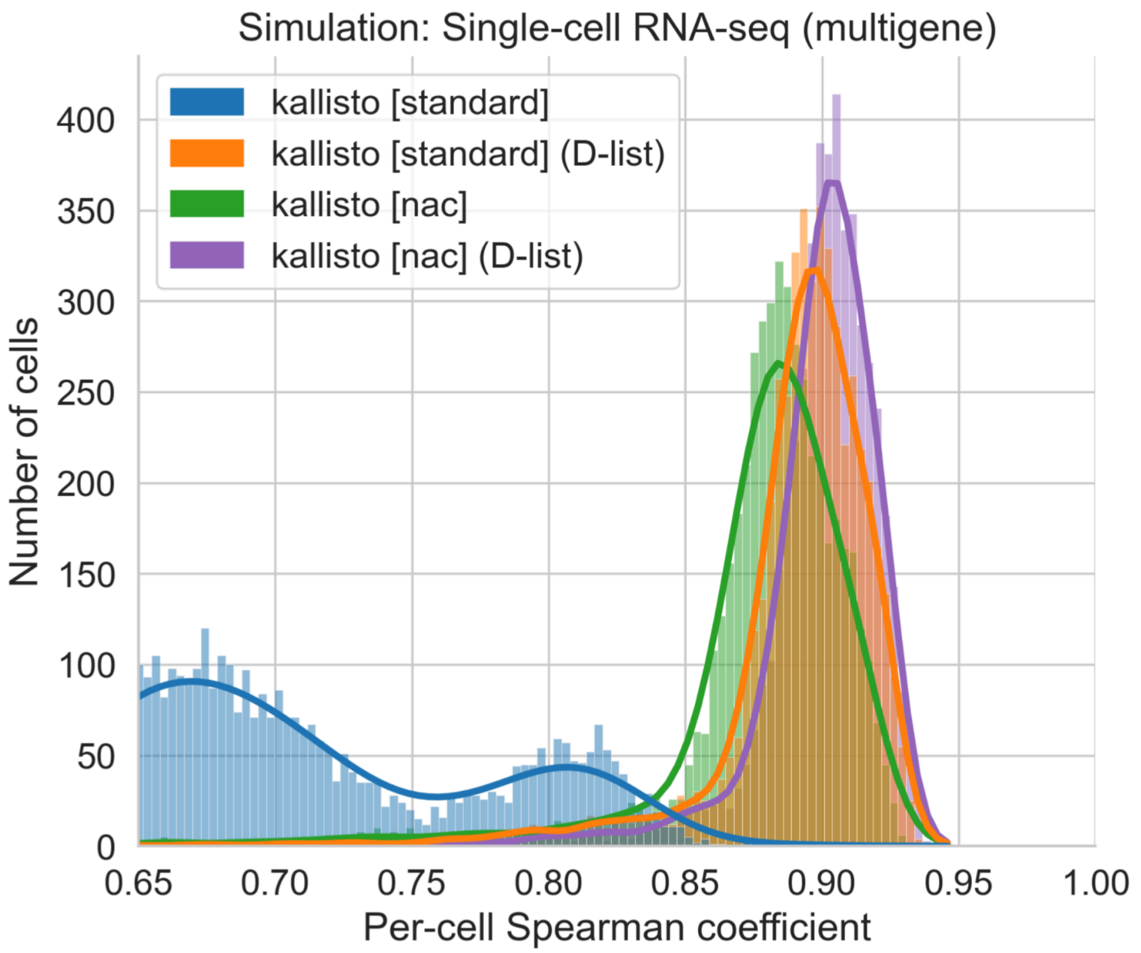


| Index type | D-list | mult | Median ρ* | Median r | RMSE | FPR | FNR |
|---|---|---|---|---|---|---|---|
| standard | | | 0.679939 | 0.991087 | 0.682919 | 0.019123 | 0.002950 |
| standard | ✓ | | 0.896340 | 0.997867 | 0.287012 | 0.000783 | 0.003218 |
| nac | | | 0.883905 | 0.994031 | 0.555755 | 0.000626 | 0.003342 |
| nac | ✓ | | 0.902738 | 0.997835 | 0.289252 | 0.000197 | 0.003367 |

**Supplementary Figure 3**: Assessment of kallisto quantification results with multimapping mode enabled on simulated single-cell RNA-seq reads that do not include multi-gene reads.

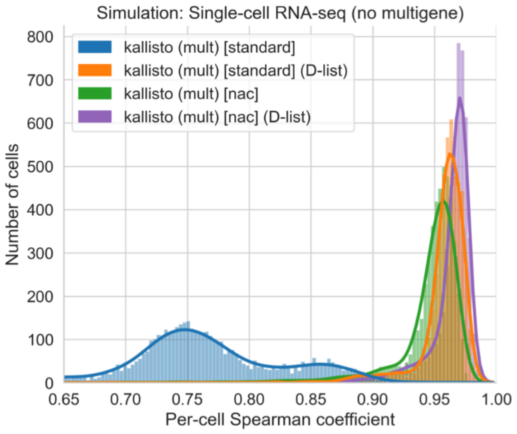


| Index type | D-list | mult | Median ρ* | Median r | RMSE | FPR | FNR |
|---|---|---|---|---|---|---|---|
| standard | | ✓ | 0.756161 | 0.992337 | 0.653089 | 0.050188 | 0.000241 |
| standard | ✓ | ✓ | 0.961758 | 0.999728 | 0.104715 | 0.002396 | 0.000431 |
| nac | | ✓ | 0.953292 | 0.995827 | 0.506452 | 0.023166 | 0.000517 |
| nac | ✓ | ✓ | 0.968557 | 0.999734 | 0.105673 | 0.018768 | 0.000542 |

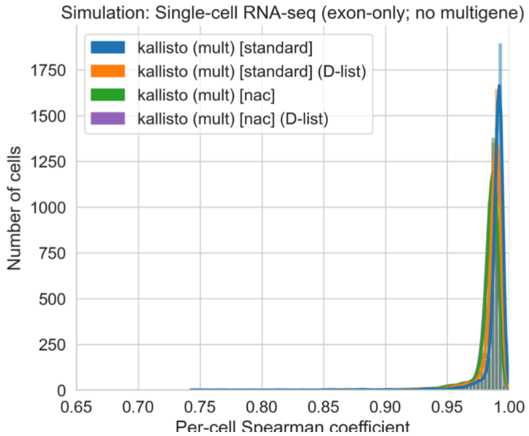


| Index type | D-list | mult | Median ρ* | Median r | RMSE | FPR | FNR |
|---|---|---|---|---|---|---|---|
| standard | | ✓ | 0.991460 | 0.999955 | 0.039947 | 0.000249 | 0.000321 |
| standard | ✓ | ✓ | 0.989052 | 0.999925 | 0.050895 | 0.000173 | 0.000448 |
| nac | | ✓ | 0.986388 | 0.999891 | 0.061643 | 0.000508 | 0.000554 |
| nac | ✓ | ✓ | 0.985920 | 0.999878 | 0.065441 | 0.000473 | 0.000575 |

**Supplementary Figure 4:** The different count matrices and their combinations produced from the mouse embryo Visium CytAssist 11mm FFPE spatial transcriptomics dataset with 832,193,962 reads.

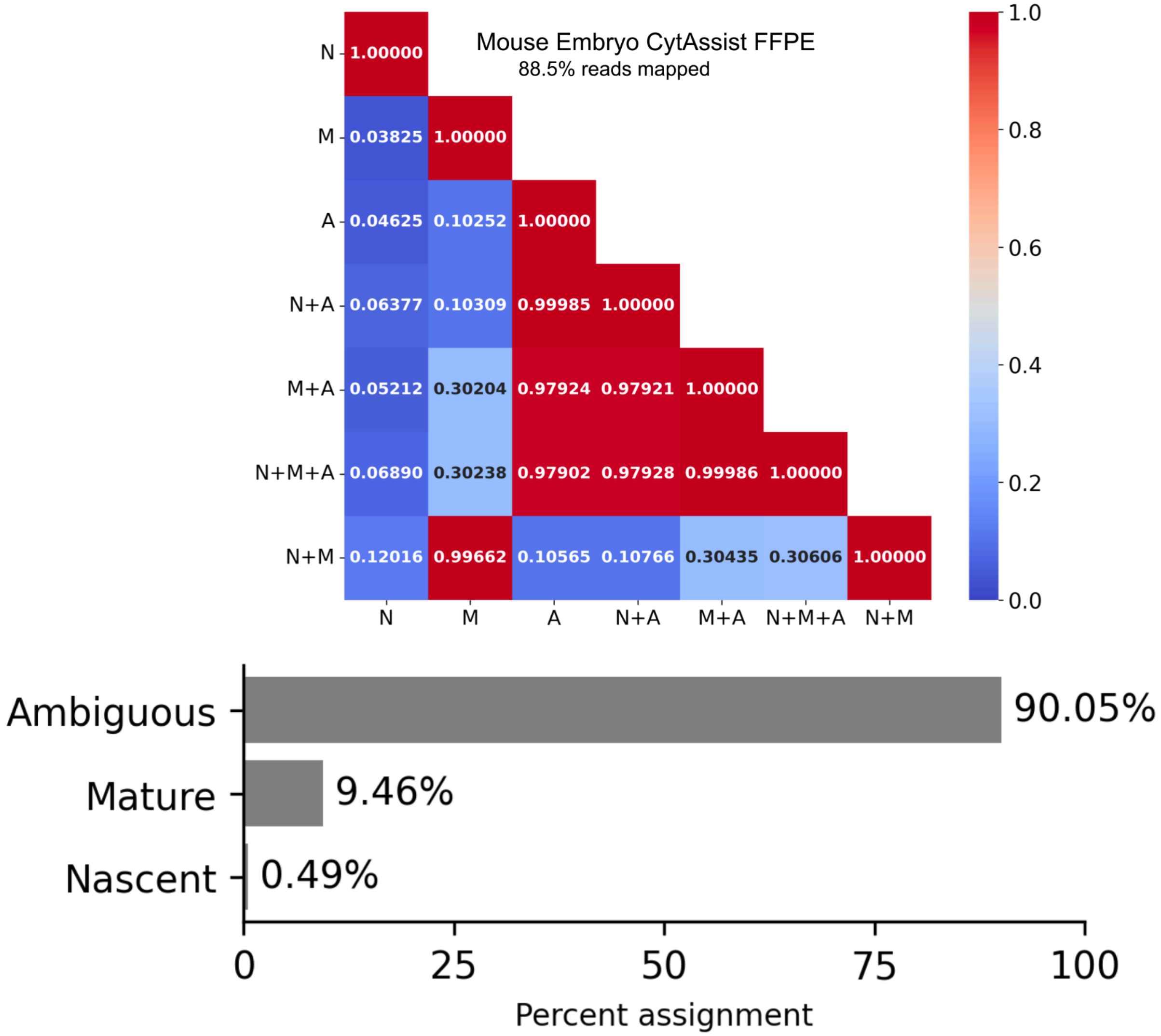

**Supplementary Figure 5**: Depiction of the distinct count matrices that can be studied with data from a single-cell or single-nucleus genomics experiment.

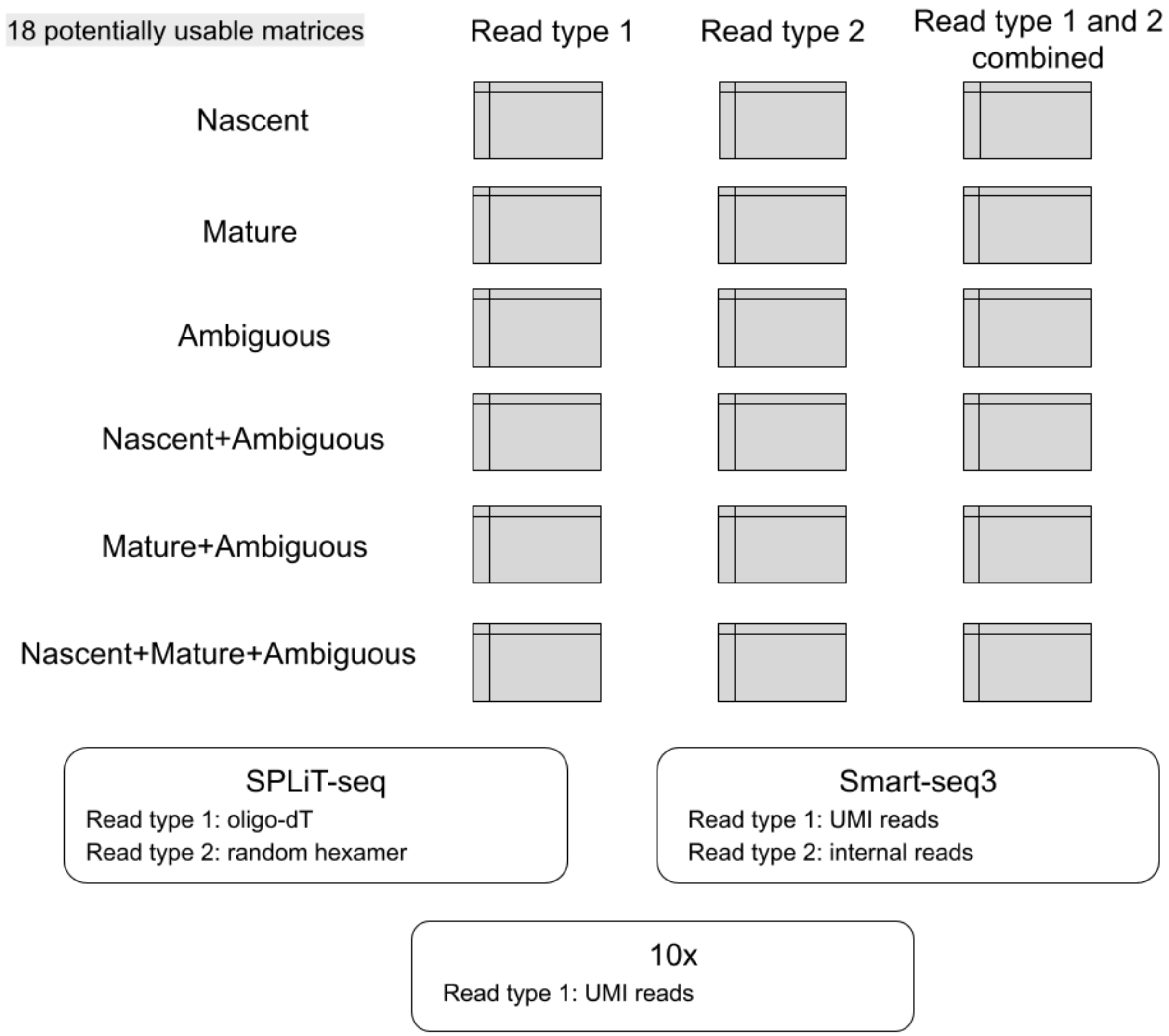

**Supplementary Figure 6**: Performance comparison of different implementations of the nac index type.

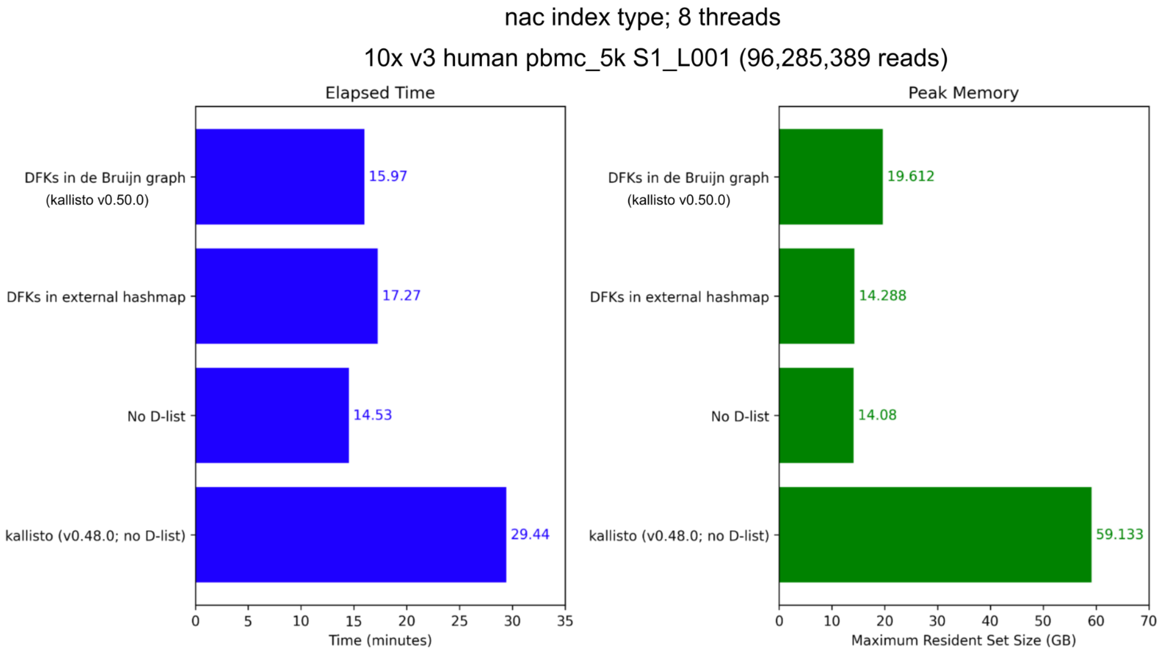

**Supplementary Table 1**: Evaluation metrics of kallisto on simulated data as a function of DFK overhang length.

| Index type | Overhang | mult | Median ρ* | Median r | RMSE | FPR | FNR | *k*-mers | DFKs |
|---|---|---|---|---|---|---|---|---|---|
| Simulation: Single-cell RNA-seq (no multigene) | | | | | | | | | |
| standard | 2 | | 0.973984 | 0.999819 | 0.080112 | 0.000739 | 0.000495 | 113,209,587 | 9,649,025 |
| standard | 3 | | 0.974748 | 0.999824 | 0.078881 | 0.000685 | 0.000506 | 113,209,587 | 15,696,903 |
| standard | 4 | | 0.97535 | 0.999826 | 0.07816 | 0.000643 | 0.000513 | 113,209,587 | 22,293,140 |
| standard | 5 | | 0.975842 | 0.99983 | 0.077472 | 0.000608 | 0.000519 | 113,209,587 | 29,278,838 |
| standard | 6 | | 0.976265 | 0.999833 | 0.076833 | 0.00058 | 0.000524 | 113,209,587 | 36,550,263 |
| nac | 2 | | 0.980336 | 0.999812 | 0.083905 | 0.000202 | 0.000618 | 1,398,470,117 | 21,338,179 |
| nac | 3 | | 0.980491 | 0.999814 | 0.08297 | 0.000197 | 0.000619 | 1,398,470,117 | 30,833,829 |
| nac | 4 | | 0.980585 | 0.999816 | 0.082474 | 0.000193 | 0.00062 | 1,398,470,117 | 39,760,674 |
| nac | 5 | | 0.980698 | 0.999817 | 0.082168 | 0.000189 | 0.000621 | 1,398,470,117 | 48,233,865 |
| nac | 6 | | 0.980771 | 0.999819 | 0.081806 | 0.000185 | 0.000622 | 1,398,470,117 | 56,344,346 |
| mult: the multimapping quantification mode is enabled (not enabled for any of the runs here). Overhang: The number of *k*-mers flanking a unitig to be considered a DFK. ρ*: Modified spearman correlation. r: Pearson correlation. RMSE: root mean squared error. FPR: false positive rate. FNR: false negative rate. See **Methods** for details. | | | | | | | | | |

**Supplementary Table 2**: Evaluation of the STARsolo, Cell Ranger, alevin-fry and cellCounts single-cell RNA-seq programs on simulated data generated using the STARsolo simulation framework.

| Program | Run mode | Median ρ* | Median r | RMSE | FPR | FNR |
|---|---|---|---|---|---|---|
| Simulation: Single-cell RNA-seq (exon-only; no multigene) | | | | | | |
| STARsolo | | 0.993564 | 0.999968 | 0.030486 | 0.000053 | 0.000270 |
| Cell Ranger | --include-introns=false | 0.956866 | 0.999547 | 0.232195 | 0.000030 | 0.000343 |
| alevin-fry | splici align | 0.984427 | 0.999902 | 0.069541 | 0.000030 | 0.000709 |
| alevin-fry | splici sketch | 0.982900 | 0.999889 | 0.062949 | 0.000031 | 0.000743 |
| alevin-fry | align | 0.990880 | 0.999931 | 0.060543 | 0.000030 | 0.000466 |
| alevin-fry | sketch | 0.991528 | 0.999948 | 0.043601 | 0.000031 | 0.000393 |
| cellCounts | | 0.915796 | 0.994174 | 0.521436 | 0.000239 | 0.001337 |
| Simulation: Single-cell RNA-seq (no multigene) | | | | | | |
| STARsolo | | 0.991877 | 0.999940 | 0.042743 | 0.000140 | 0.000270 |
| Cell Ranger | --include-introns=false | 0.955377 | 0.999499 | 0.232933 | 0.000081 | 0.000343 |
| alevin-fry | splici align | 0.971626 | 0.998672 | 0.288308 | 0.000314 | 0.000702 |
| alevin-fry | splici sketch | 0.960025 | 0.997419 | 0.414070 | 0.000764 | 0.000739 |
| alevin-fry | align | 0.879684 | 0.998145 | 0.312199 | 0.005676 | 0.000446 |
| alevin-fry | sketch | 0.778933 | 0.995372 | 0.527728 | 0.016063 | 0.000357 |
| cellCounts | | 0.825302 | 0.993000 | 0.532384 | 0.005890 | 0.001299 |

Modified spearman correlation. r: Pearson correlation. RMSE: root mean squared error. FPR: false positive rate. FNR: false negative rate. See **Methods** for details.

alevin-fry options:

- splici align: Enabling the index used by alevin-fry that contains introns as well as selective alignment mode.
- splici sketch: Enabling the index used by alevin-fry that contains introns without selective alignment mode.
- align: Selective alignment enabled and index is a standard transcriptome index that does not include introns in alevin-fry.
- sketch: Selective alignment disabled and index is a standard transcriptome index that does not include introns in alevin-fry.

For Cell Ranger, version 7 was used with the include-introns option set to false in order to mimic the default behavior of versions 1,2,3,4,5 and 6.

**Supplementary Table 3**: Evaluation metrics of the kallisto, alevin-fry, and STARsolo single-cell RNA-seq programs on simulated data generated using the STARsolo simulation framework with errors introduced into sequencing reads.

| Mismatches | Deletions | Insertions | Program | Run mode | Median ρ* | Median r | RMSE | FPR | FNR |
|---|---|---|---|---|---|---|---|---|---|
| Simulation: Single-cell RNA-seq (no multigene) | | | | | | | | | |
| 1 | 1 | 1 | kallisto | standard (no D-list) | 0.629277 | 0.993046 | 0.575156 | 0.030304 | 0.001565 |
| 1 | 1 | 1 | kallisto | standard (D-list) | 0.87452 | 0.998388 | 0.276873 | 0.003689 | 0.002376 |
| 1 | 1 | 1 | kallisto | nac (no D-list) | 0.890008 | 0.995764 | 0.484241 | 0.001393 | 0.002872 |
| 1 | 1 | 1 | kallisto | nac (D-list) | 0.900935 | 0.998259 | 0.29544 | 0.001023 | 0.002973 |
| 1 | 1 | 1 | alevin-fry | splici align | 0.673962 | 0.989232 | 1.592164 | 0.000077 | 0.018216 |
| 1 | 1 | 1 | alevin-fry | splici sketch | 0.899393 | 0.99736 | 0.39301 | 0.001391 | 0.002823 |
| 1 | 1 | 1 | STARsolo | | 0.911674 | 0.998962 | 0.271059 | 0.000888 | 0.002918 |
| 4 | 0 | 0 | kallisto | standard (no D-list) | 0.612826 | 0.992954 | 0.570622 | 0.030607 | 0.002184 |
| 4 | 0 | 0 | kallisto | standard (D-list) | 0.85271 | 0.998136 | 0.305361 | 0.004151 | 0.002991 |
| 4 | 0 | 0 | kallisto | nac (no D-list) | 0.870739 | 0.995728 | 0.483343 | 0.001446 | 0.003621 |
| 4 | 0 | 0 | kallisto | nac (D-list) | 0.881683 | 0.997985 | 0.330969 | 0.001098 | 0.003706 |
| 4 | 0 | 0 | alevin-fry | splici align | 0.819814 | 0.996343 | 0.76258 | 0.000141 | 0.008301 |
| 4 | 0 | 0 | alevin-fry | splici sketch | 0.878774 | 0.9971 | 0.415686 | 0.00136 | 0.003727 |
| 4 | 0 | 0 | STARsolo | | 0.948253 | 0.999501 | 0.141767 | 0.000947 | 0.001244 |
| 0 | 2 | 2 | kallisto | standard (no D-list) | 0.586625 | 0.993183 | 0.557911 | 0.030393 | 0.002935 |
| 0 | 2 | 2 | kallisto | standard (D-list) | 0.831982 | 0.997594 | 0.381203 | 0.00433 | 0.003804 |
| 0 | 2 | 2 | kallisto | nac (no D-list) | 0.851074 | 0.99581 | 0.500121 | 0.001891 | 0.004287 |
| 0 | 2 | 2 | kallisto | nac (D-list) | 0.861993 | 0.997502 | 0.403226 | 0.001458 | 0.004384 |
| 0 | 2 | 2 | alevin-fry | splici align | 0.412969 | 0.924791 | 3.764902 | 0.000033 | 0.053289 |
| 0 | 2 | 2 | alevin-fry | splici sketch | 0.852768 | 0.996873 | 0.466828 | 0.00178 | 0.004671 |
| 0 | 2 | 2 | STARsolo | | 0.763642 | 0.994432 | 0.948656 | 0.000552 | 0.010949 |

Modified spearman correlation. r: Pearson correlation. RMSE: root mean squared error.
FPR: false positive rate. FNR: false negative rate. See **Methods** for details.
alevin-fry options:

- splici align: Enabling the index used by alevin-fry that contains introns as well as selective alignment mode.
- splici sketch: Enabling the index used by alevin-fry that contains introns without selective alignment mode.

**Supplementary Text 1**

Response to: He D, Soneson C, Patro R. "Understanding and evaluating ambiguity in single-cell and single-nucleus RNA-sequencing". Biorxiv. 2023. https://doi.org/10.1101/2023.01.04.522742

D. He, C. Soneson and R. Patro wrote a reply to an initial version of this preprint, to which we provide a point-by-point response below:

“However, we find that it leaves a large fraction of reads classified as ambiguous, and, in practice, allocates these ambiguous reads in an all-or-nothing manner, and differently between single-cell and single-nucleus RNA-seq data.”

He et al. correctly state that classifying reads as mature, nascent, or ambiguous results in a large fraction of reads being classified as ambiguous. We agree that the presence of a substantial number of ambiguous reads is an important consideration, and as He et al. state later on, this is something we "acknowledged in [our] manuscript itself”. However, the ambiguous reads are an integral part of analysis, and we do not suggest discarding them, believing that they can provide valuable insights in many contexts.

He et al. state that we allocate ambiguous reads in an “all-or-nothing manner”. This “all-or-nothing manner” resolution has been, and continues to be, true for the field for the past several years where reads are always assigned to one splicing status or another. This approach to handling ambiguous reads is not something unique to, or originating with, our work. Rather, our work presents an approach to accurately and correctly identify ambiguity where it exists.

He et al. make a valid point that ambiguous reads may need to be allocated differently by different assay types; in some assays, an ambiguous read might be more likely to have come from a nascent transcript; in other assays, an ambiguous read might be more likely to have come from a mature transcript. We provide an approach to allocate ambiguous reads accordingly, providing a framework for us and others to evaluate how modeling splicing in single-cell assays might differ from modeling splicing in single-nucleus assays. We concur with He et al., that the nascent/mature/ambiguous “proportions can vary from sample to sample, cell type to cell type, across genes, and, of course between different assays.”

“While the conservative motivation of this classification scheme makes sense conceptually, its current practical utility is questionable”

He et al. consider our work “questionable”. However, the classification has proven to be extremely useful for us and for others. It has allowed us to conduct detailed analyses, including examining coverage over exons, introns, and splice junctions across different assays and cell types. Such analysis, which is crucial for quality control and assay comparison, has also been utilized by He et al. in Table 1 of their paper. Their findings, which indicate substantial variation in proportions between different assays, underscore the practicality of our approach. Furthermore, the A-SNR model proposed by He et al. incorporates our method, which we believe further affirms its utility. Examining the nascent/mature/ambiguous quantifications separately and analyzing what parameters might differ between the three is the premise for some exciting ongoing work in our lab.

“the UMIs falling into the ambiguous class are allocated differently and precisely in the manner that will most benefit the proposed method in the assessment being carried out”

We appreciate the concerns by He et al., however, this was the purpose of a simulation. In some assays, an ambiguous read might be more likely to have originated from a nascent transcript; in other assays, an ambiguous read might be more likely to have originated from a mature transcript. Ideally ambiguous reads are allocated in a manner most consistent with the assay, therefore providing the most benefit when modeling the biology. The simulation is, of course, a purely hypothetical situation but illustrates an instance where we may want to tune our allocation accordingly when we don’t have a solution to the nascent/mature identifiability problem.

“However, the authors do not consistently apply this definition in their manuscript.”

See previous points. Additionally, in the first version of our preprint, we had already acknowledged the treatment of ambiguous reads as “unsatisfactory” in the discussion section, and discussed avenues for future research (i.e. probabilistic classification), which was expanded upon by He et al. in their reply.

“Also, it is worth noting that the classification scheme of alevin-fry is a result of the way the splici index is constructed, and is adopted to conform to prevailing convention”

Similarly, in our own work in kallisto | bustools, we had constructed its index differently in prior versions, in an approach that resulted in different classification schemes when producing spliced/unspliced matrices. Upon the need to examine ambiguity, we have updated our index (as have the developers of alevin-fry done).

"Simply classifying these UMIs as ambiguous, while a conservative approach, vastly reduces the amount of usable data under existing downstream processing pipelines, further exacerbating the challenges posed by the sparse sampling of the most prevalent single-cell sequencing technologies."

It is important to again clarify that we do not advocate for the discarding of ambiguous UMIs. On the contrary, we believe that identifying UMIs as ambiguous adds a layer of valuable information to our data analysis. As noted by He et al., throughout our manuscript, we have often been allocating ambiguous UMIs to nascent or mature molecules. As such, we are not only preserving but enhancing the utility of the data by providing a more nuanced understanding of its nature.

"Further, assigning them entirely to either a spliced or unspliced status, depending on the type of assay being processed, does little to ameliorate the problem beyond prevailing convention."

It's true, as He et al. have noted, and as our revised analysis confirms, that for certain types of analyses such as single-nucleus RNA-seq quantification, the difference in outcomes between assigning ambiguous UMIs to the 'nascent' status, versus other approach such as summing all UMIS, may lead to similar coarse grained results. However, it's important to emphasize that this is just one particular type of analysis. There are other analytical scenarios, particularly those involving the modeling of splicing, where the allocation of UMIs to a specific status becomes a critical factor. We are continually exploring these different analytical dimensions to enrich our understanding and application of RNA-seq data.

"Difficulty in reproducing certain results due to absent files and code"

Our lab routinely posts code to facilitate reproducing results. We have updated our repository with code available to reproduce the newer results in this manuscript. We welcome the posting of GitHub issues if there are any problems running the code or if a file is missing.

“Surprisingly, the approach proposed by Hjörleifsson and Sullivan et al. does not actually categorize the splicing status of each UMI.”

As could be readily inferred through the referenced GitHub links, the new version of kallisto was in active development and had not been distributed via an official release. We posted the preprint to obtain feedback from the community on what could be improved. Much of the feedback was extremely valuable, substantially improved kallisto, and spurred excellent ideas for new projects that we are enthusiastically pursuing. Much of the current functionalities of kallisto/bustools/kb-python were only possible thanks to the very helpful feedback we received on this preprint. We had been actively working on the “splicing status' ' categorization and indexing of the nascent transcripts at the time (as could be discerned from our GitHub commits and comments). It’s important to note that categorizing the “splicing status” (akin to the alevin-fry spliceu index) required modifications to three additional tools: bustools, kb-python, and ngs-tools, not just kallisto. The preprint was about the new functionalities of kallisto only and therefore focused on production of a single count matrix at a time. We are pleased to report that, since then, we have released stable versions of both kallisto and bustools that can handle the addition of nascent transcripts in the index.

“Discarding — rather than resolving and quantifying — the fragments that arise from the unexpected (or less dominant) splicing status in a sample may result in misallocation of UMIs that could have been properly recovered”

1. To produce three count matrices (which allow proper allocation of UMIs into splicing statuses), the new version of kb-python (that had been a work in progress at the time the original preprint was posted) is now in its stable release and can now easily do so.
2. kallisto now has a --dfk-onlist option that results in DFK-containing reads being preserved (with a distinct element added to its equivalence class) rather than being discarded. In some of our recent workflows that make heavy use of D-list masking, this option is always enabled and has proven to be extremely valuable to identify whether a specific pseudoalignment crossed a DFK.

“Unfortunately, since STARsolo aligns the reads in a (potentially) spliced fashion to the genome, the same simple strategy cannot be used to easily assign an inferred status to each read from the results produced by STARsolo.”

It is indeed unfortunate that STARsolo cannot easily assign splicing status to each read – this highlights one advantage of alternative tools over STARsolo when performing this type of analysis.

“This difference appears to be due, almost entirely, to the fact that the generate_cDNA+introns.py script does not reverse complement nascent transcripts arising from genes on the negative strand prior to indexing”

This was likely the case and we thank He et al. for bringing this to our attention. In this revised manuscript, we have corrected the single-nucleus N+A vs. N+A+M analyses.
